# Uropathogenic *Escherichia coli* wield enterobactin-derived catabolites as siderophores

**DOI:** 10.1101/2023.07.25.550588

**Authors:** Zongsen Zou, John I. Robinson, Lindsey K. Steinberg, Jeffrey P. Henderson

## Abstract

Uropathogenic *E. coli* (UPEC) secrete multiple siderophore types to scavenge extracellular iron(III) ions during clinical urinary tract infections, despite the metabolic costs of biosynthesis. Here we find the siderophore enterobactin and its related products to be prominent components of the iron-responsive extracellular metabolome of a model UPEC strain. Using defined enterobactin biosynthesis and import mutants, we identify lower molecular weight, dimeric exometabolites as products of incomplete siderophore catabolism, rather than prematurely released biosynthetic intermediates. In *E. coli,* iron acquisition from iron(III)-enterobactin complexes requires intracellular esterases that hydrolyze the siderophore. Although UPEC are equipped to consume the products of completely hydrolyzed enterobactin, we find that enterobactin and its derivatives may be incompletely hydrolyzed to yield products with retained siderophore activity. These results are consistent with catabolic inefficiency as means to obtain more than one iron ion per siderophore molecule. This is compatible with an evolved UPEC strategy to maximize the nutritional returns from metabolic investments in siderophore biosynthesis.

## INTRODUCTION

Urinary tract infections (UTI) are among the most common outpatient and inpatient infections encountered by physicians (1–4). *Escherichia coli* is the bacterial species most commonly associated with UTI, accounting for about 70% ∼ 95% of clinical cases (5,6). Clinical *E. coli* isolates associated with UTI that exhibit polymorphisms in conserved genes (7–9) and carry accessory genes associated with increased pathogenic potential are designated as uropathogenic *E. coli* (UPEC) (2,4,10). Prominent among these virulence-associated adaptions are iron uptake systems, such as siderophores, which use distinctive chemical groups to competitively bind iron and render it selectively bioavailable to support bacterial growth (2,3,10–14). In UPEC, siderophore iron-acquisition systems have been identified as both colonization factors and virulence factors during UTI pathogenesis (15–19). The enterobactin, salmochelin, yersiniabactin, and aerobactin siderophore systems have all been associated with *E. coli* strains that causing extraintestinal infections (20–22).

Siderophores are specialized, secreted metabolites (exometabolites) that are synthesized by non-essential bacterial pathways and competitively chelate extracellular iron(III) during the iron-limited growth conditions characteristic of infection microenvironments (16,17,23,24). The resulting iron(III)-siderophore complexes are selectively imported by bacterial transporters as an iron source. *E. coli* and many other gram-negative bacteria actively transport iron-siderophore complexes through outer membrane receptors using the cytoplasmic membrane□localized TonB/ExbB/ExbD complex, which transduces energy from the proton motive force (25–27). Siderophore biosynthesis and transport systems are regulated by the ferric uptake regulator (Fur), a transcriptional repressor that downregulates siderophore gene transcription in conditions associated with high cytosolic iron (28).

All UPEC carry the conserved enterobactin system and may encode up to three additional siderophore systems, each associated with chemically distinctive exometabolomes (29–31). Biosynthesis of these additional exometabolites incurs additional metabolic demands (32), suggesting that their sustained presence in clinical populations is associated with siderophore-specific payoffs. For example, the salmochelin system, encoded by genes in the *iroA* cassette, glucosylates enterobactin to improve its aqueous solubility and evade sequestration by the host immune protein lipocalin-2/siderocalin/NGAL (14,33,34). The yersiniabactin system in UPEC supports multiple non-siderophore functions not associated with enterobactin or salmochelin (35,36). Yersiniabactin production incurs metabolic costs, which appear to be mitigated by an ability to recycle the intact siderophore to support multiple rounds of metal ion import (37) and an additional quorum-sensing regulatory input that emphasizes biosynthesis in diffusionally restricted or crowded environments where the siderophore is more likely to remain nearby (38).

Enterobactin (Ent) is detectable in the urine of patients with urinary tract infections, where its synthesis is required to evade growth inhibition by Lipocalin-2 (13,14). Ent achieves exceptional iron (III) affinity (K_d_ ≈ 10^-52^ M) with three catechol (1,2-dihydroxybenzene) groups that provide all six coordination sites for iron(III) (10,39). Ent is synthesized by a nonribosomal peptide synthetase (NRPS) system encoded by *entABCDEF*. This NRPS system is a molecular assembly line that synthetizes Ent by repeatedly forming enzyme-bound *N*-(2,3-dihydroxybenzoyl)serine (DHBS) and linking them via ester bonds until a cyclic trilactone core composed of three DHBS is released (40,41). In UPEC expressing the *iroA* cassette, the glucosyltransferase IroB further modifies Ent catechols with up to three distinctive C-linked glucoses (10,42). Iron retrieval from imported iron(III) Ent complexes (with or without C-glucose modifications) requires dissociation through both esterase-catalyzed Ent hydrolysis (by Fes and/or IroD) and iron (III) reduction to iron(II) (43–45).

In this study we examined the enterobactin biosynthetic pathway’s contribution to the iron-dependent UPEC exometabolome. We measured used targeted mutant strains and chemical complementation with purified products to assess the catabolic origins of short-length catechol exometabolites. To assess the nutritional potential of siderophore catabolism, we used reverse stable isotope labeling to find that 2,3-dihydroxybenoic acid from outside the biosynthetic pathway could be used for Ent biosynthesis. Finally, we used a siderophore-dependent growth condition to evaluate the siderophore potential of non-trimeric enterobactin metabolites found in the UPEC exometabolome. Our findings are consistent with a catabolic network that has evolved to maximize the iron delivery potential of Ent biosynthesis.

## RESULTS

### Enterobactin and the iron-responsive exometabolome in uropathogenic E. coli

To define the iron-responsive exometabolome of uropathogenic *E. coli* and its relationship to the *ent-*encoded biosynthetic pathway, we compared small molecule profiles in conditioned media from the model UPEC strain UTI89 and its isogenic biosynthesis-deficient mutant, UTI89Δ*entB* (21), in low and high iron conditions (32) using LC-MS. Sparse principle components analysis (sPCA) of these data demonstrated discrete groupings along principle component 1 (PC1, **Fig. 1A**) for the experimental groups that account for 26.8% of total variation between specimens (**Fig. 1A, supplemental Fig. S1A**). The greatest separation was evident between wild type UTI89 grown in low iron medium and the other experimental groups, which were fully separated along PC1. Logistic regression of PC1 values to classify these two PC1 exometabolome clusters yielded a prediction accuracy of 1.0 (SD = 0, **supplemental Fig. S1B**) and an AUC of 1.0 (SD = 0, **supplemental Fig. S1C**) with four-fold cross validation. PC1 differences did not correspond to inter-group differences in growth density. **(supplemental Fig. S2)**. The distinctive PCA grouping of wild type UTI89 grown in low iron corresponds with detection of enterobactin, the canonical, eponymous product of the enterobactin biosynthesis pathway (**Fig. 1B**). Together, these results are consistent with a prominent role for the enterobactin biosynthetic pathway in defining the iron-responsive UTI89 exometabolome.

**Figure 1.**
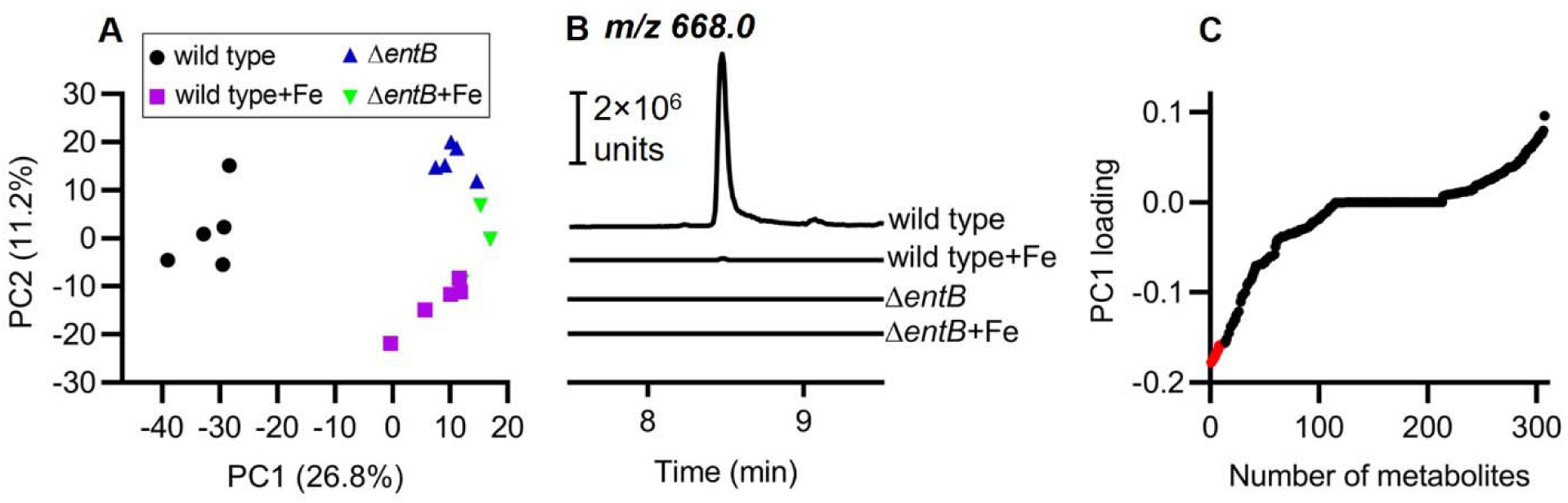
The enterobactin biosynthetic pathway is a prominent contributor to the iron-responsive uropathogenic *E. coli* UTI89 exometabolome. **(A)** Score plot from sparse PCA (sPCA) analysis of media conditioned by UTI89 grown in low and high iron media (wild type and wild type+Fe, respectively) and the enterobactin-null mutant UTI89Δ*entB* in low and high iron media (*entB* and *entB* +Fe, respectively). High iron medium is achieved by addition of 100 µM FeCl_3_ **(B)** LC-MS/MS chromatograms corresponding to the precursor-product ions from enterobactin for each experimental group. Chromatograms are displayed in identical ion current unit scales. **(C)** PC1-loadings plot demonstrates that multiple ions contribute to PC1. The top 13 metabolites with greater abundance in wild type UTI89 (lower PC1 value) are identified as red data points.

### Multiple enterobactin-associated products define the UTI89 exometabolome

The exometabolites that define principal components in these data may be identified by loadings analysis. Loadings analysis of PC1, corresponding to the greatest variance in exometabolites, identified multiple molecules enterobactin alone (**Fig. 1C**). Detailed mass spectrometric and chromatographic analyses of the 13 molecular features with the largest PC1 loadings associated with the UTI89 exometabolome under iron-restricted conditions (**supplemental Fig. S3-12**) identified a series of 10 *N*-(2,3-dihydroxybenzoyl)serine (DHBS) polymers (**Fig. 2A**) consistent with enterobactin (Ent) and salmochelin biosynthesis (46–48) (**Table S1**). These include cyclic and linear DHBS trimers with 0, 1, or 2 C-glucosylations, DHBS dimers with 0, 1, or 2 C-glucosylations, and monomeric DHBS previously reported in an avian pathogenic *E. coli* (APEC) strain. Unlike the APEC strain, UTI89 did not produce tri-glucosylated Ent products (54), consistent with inter-strain differences in the Ent exometabolome that are not explained by *iroA* alone. To more precisely quantify these exometabolites, we constructed a high-resolution targeted LC-MS/MS multiplexed selected reaction monitoring (LC-MRM) method (**Table 1**). We confirmed that all 10 products were present in low iron media conditioned by wild type UTI89, were significantly diminished in high iron media conditioned by UTI89, and were undetectable in any media conditioned by UTI89Δ*entB* (**Fig. 2B, supplemental Fig. S13, supplemental Fig. S14**, *P* < 0.001). In an *iroA-*null strain (UTI89Δ*ybtS*Δ*iroA*) that lacks the C-glucosylation pathway, C-glucosylated exometabolites were absent, while non-glucosylated exometabolites were elevated, (**Fig. 2B, supplemental Fig. S14**, *P* < 0.001), consistent with the precursor-product relationship between these exometabolites. Together, these results connect iron-associated biosynthetic activity in uropathogenic *E. coli* to multiple enterobactin-related exometabolites extending beyond the canonical trimeric DHBS products.

**Figure 2.**
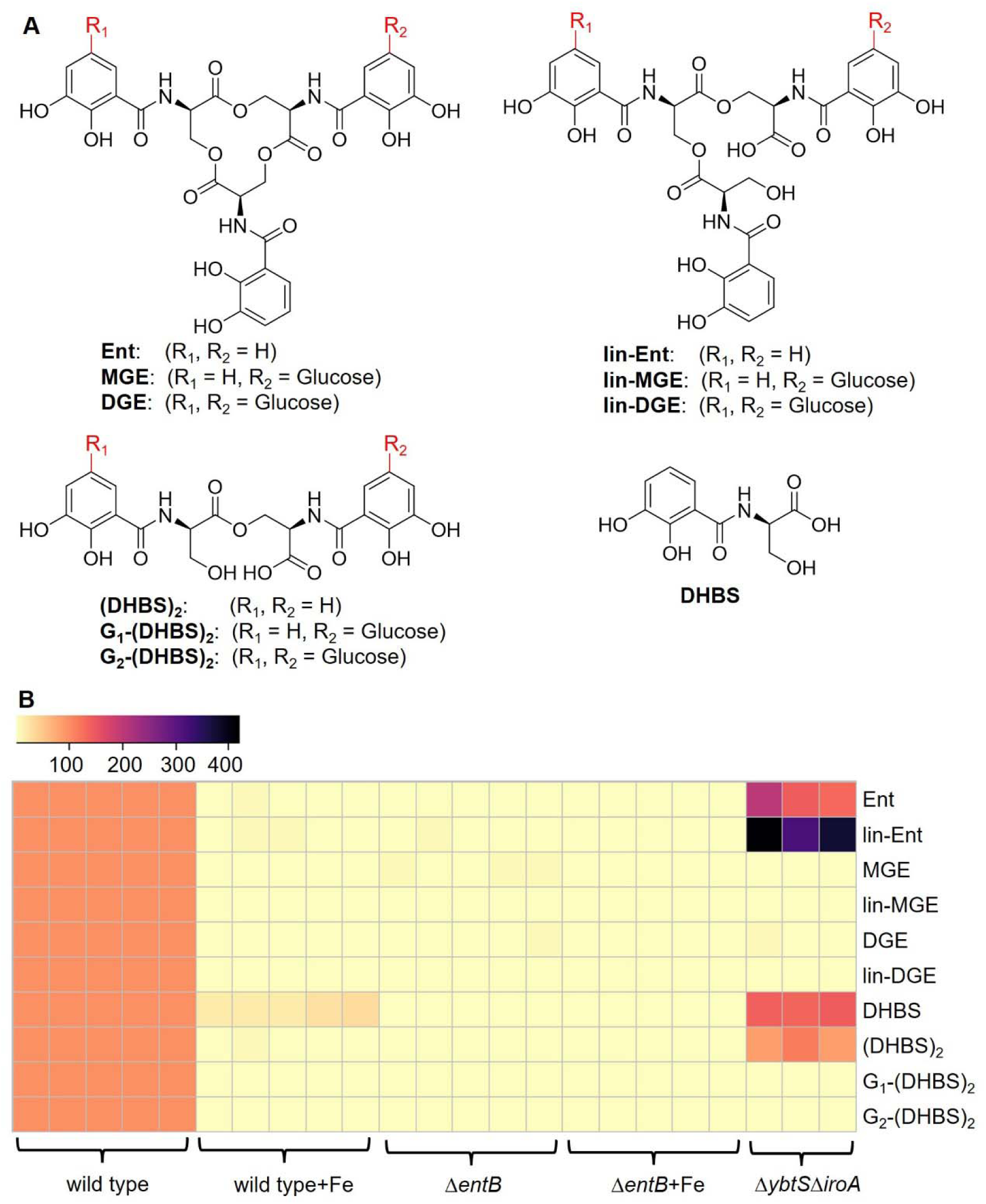
Exometabolites associated with the iron-responsive UTI89 exometabolome. **(A)** Chemical structures of the ten enterobactin*-*associated exometabolites identified by comparative metabolomic analysis, including enterobactin (Ent), monoglucosylated enterobactin (MGE), diglucosyalted enterobactin (DGE), linear enterobactin (lin-Ent), linear monoglucosylated enterobactin (lin-MGE), linear diglucosylated enterobactin (lin-DGE), *N*-(2,3-dihydroxybenzoyl)serine dimer [(DHBS)_2_], monoglucosylated *N*-(2,3-dihydroxybenzoyl)serine dimer [G_1_-(DHBS)_2_], diglucosyalted *N*-(2,3-dihydroxybenzoyl)serine dimer [G_2_-(DHBS)_2_], and *N*-(2,3-dihydroxybenzoyl)serine monomer (DHBS). The positions of C-glucosylated DHBS units within linear polymers have not been definitively identified. **(B)** Heatmap showing enterbactin-associated exometabolite concentrations in media with iron supplementation or defined biosynthetic mutants of UTI89. Intensity represents concentration expressed as ratio of LC-MS/MS peak area to that of internal standard. Individual biological replicates are shown for each condition.

**Table 1.** Targeted LC-MS/MS protocols for detecting and quantifying ten *ent* siderophores and short-length products.

| Identity | Multiple reaction |  | RT (min) | Reference |
| --- | --- | --- | --- | --- |
|  | Q1 | Q3 |  |  |
| Ent | 668 | 222 | 8.47 | (21,46,49-52) |
| lin-Ent | 686 | 222 | 7.21 | (21,46,47,49) |
| MGE | 832 | 222 | 6.94 | (46,47,49) |
| lin-MGE | 848 | 222 | 5.89 | (46,47,49) |
| DGE | 993 | 222 | 5.95 | (46,47,49,53) |
| lin-DGE | 1010 | 222 | 4.98 | (21,46,47,49,53) |
| (DHBS)<sub>2</sub> | 464 | 222 | 5.92 | (46,49-52) |
| G<sub>1</sub>-(DHBS)<sub>2</sub> | 625 | 222 | 4.69 | (46,47,49,53) |
| G<sub>2</sub>-(DHBS)<sub>2</sub> | 787 | 402 | 3.64 | (46,47,49) |
| DHBS | 240 | 153 | 2.34 | (46,49-52) |

### Outer membrane importers differentially affect enterobactin-associated exometabolites

While trimer products are consistent with the enterobactin biosynthetic pathway, the specific origin of short-length dimeric and monomeric products, (DHBS)_2_, (G_2_-(DHBS)_2_, G_1_-(DHBS)_2_, and DHBS is unclear. We considered that these truncated products could reflect premature release from the biosynthetic pathway (anabolic production) (11,55), spontaneous extracellular hydrolysis, or intracellular esterolysis of imported, ferric catechol siderophores (catabolic production) (46,47,49). To distinguish these possibilities, we compared UTI89 with UTI89Δ*tonB*, an isogenic mutant with a deficiency in siderophore import at the outer membrane. In *E. coli* and related gram-negative bacteria, the TonB/ExbB/ExbD complex energizes outer membrane transporters to import ferric siderophores (56). Relative to UTI89, UTI89ΔtonB cultures exhibit a strikingly dichotomous effect on enterobactin-associated exometabolites, with elevated trimer concentrations and diminished dimer and monomer concentrations (**Fig. 3**). These differences were not associated with differential growth density between groups (**supplemental Fig. S15**). These results are consistent with intracellular dimer and monomer production in events that are downstream from extracellular trimer import.

**Figure 3.**
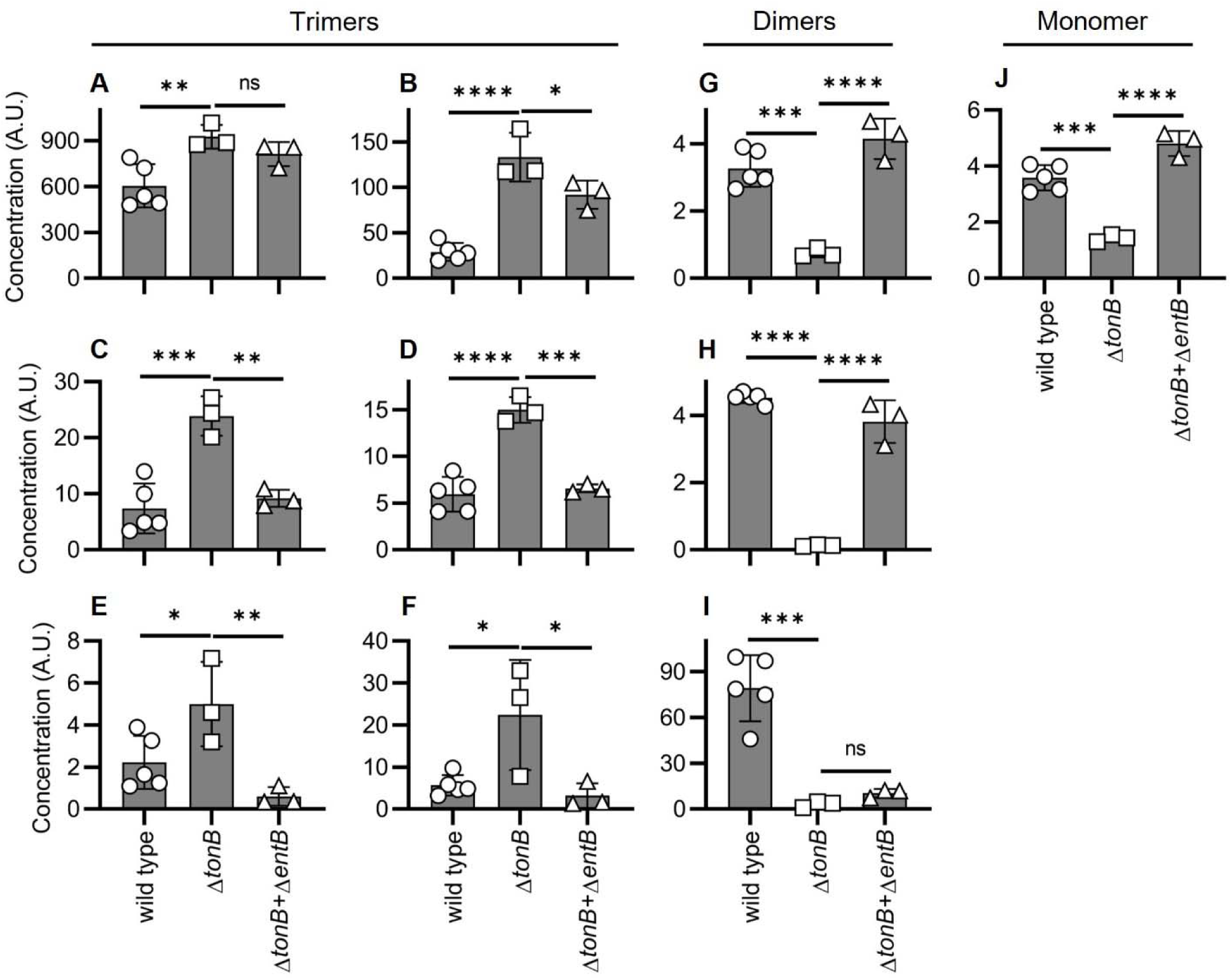
Outer membrane import differentially affects trimeric and non-trimeric enterobactin-associated exometabolites in culture. Enterobactin-associated exometabolite concentrations in media conditioned by UTI89 (wild type), an import-deficient UTI89 mutant (Δ*tonB*), or co-culture of enterobactin-null and import-deficient UTI89 mutants (Δ*tonB*+Δ*entB*). Y-axis is concentration expressed as ratio of LC-MS/MS peak area to that of internal standard. **(A)** Ent. **(B)** lin-Ent. **(C)** MGE. **(D**) lin-MGE. **(E)** DGE. **(F)** lin-DGE. **(G)** (DHBS)_2_ **(H)** G_1_-(DHBS)_2_. **(I)** G_2_-(DHBS)_2_. **(J)** DHBS. Statistics were performed using 1-way ANOVA with Dunnett’s multiple-comparison test with P ≤ 0.05 considered as statistically significant. ns: not significant. *: *P* <= 0.05. **: *P* < 0.01. ***: *P* < 0.001. ****: *P* < 0.0001.

### Co-culture with import-proficient UTI89 complements the UTI89**Δ**tonB phenotype

To further test the hypothesis that monomer and dimer exometabolites are products of siderophore catabolism, we devised a co-culture system in which UTI89Δ*tonB* is poised to serve as a siderophore-producer and enterobactin-deficient UTI89Δ*entB* as a siderophore consumer. We hypothesized that UTI89Δ*entB* import of UTI89Δ*tonB*-derived exometabolites would counteract the UTI89Δ*tonB* dimer and monomer deficiency phenotype. Compared to UTI89Δ*tonB*-conditioned media, media conditioned by the UTI89Δ*tonB* + UTI89Δ*entB* co-culture contained significantly greater monomer and dimer concentrations, and variably lower trimer concentrations (**Fig. 3**). As such, the combined *ent* exometabolome of UTI89Δ*tonB* + UTI89Δ*entB* more closely resembled that of wild type UTI89 than either mutant alone. Different levels of enterobactin-associated products were not associated with growth density differences between groups (**supplemental Fig. S15**). These results are consistent with extracellular UTI89Δ*tonB*-derived trimers as public goods that are imported by UTI89Δ*entB*, which partially catabolizes them and releases esterolysis products to the extracellular space (47,49).

### Monomer and dimer production during trimer-dependent growth

Enterobactin-associated trimers contain two or three serine-serine ester bonds and three serine-DHB peptide bonds (**Fig. 2A**) with potential for hydrolysis to yield free DHB and serine, which may become new metabolic substrates in the cytoplasm. Despite this catabolic potential, UTI89 releases incompletely hydrolyzed trimer catabolites. To determine whether this occurs during siderophore-dependent growth, we measured the enterobactin-associated exometabolomes of siderophore-null UTI89 (UTI89Δ*entB*Δ*ybtS*) cultures with trimer supplementation. Growth of this strain was rendered siderophore-dependent by addition of bovine serum albumin, a biologically relevant, non-specific binder of labile iron ions (57,58). Compared to siderophore-free controls, Ent, MGE, or DGE addition stimulated UTI89Δ*entB*Δ*ybtS* growth (**Fig. 4**) and were progressively consumed during culture (**Fig. 5A, D, G**), consistent with their canonical siderophore activity. Dimer and monomer production varied with the specific trimer provided. Ent supplementation yielded neither dimer nor monomer (**Fig. 5B, C**), MGS supplementation yielded (DHBS)_2_, G_1_-(DHBS)_2_, and DHBS (**Fig. 5E, F**), and DGS yielded G_1_-(DHBS)_2_, G_2_-(DHBS)_2_, and DHBS (**Fig. 5H, I**). Dimer C-glucosylation products are structurally consistent with the C-glucosylation structure of each trimeric substrate. These results are consistent with dimer and monomer production from esterolysis following cyclic trimer-mediated iron-delivery. The lack of dimer or monomer generation from Ent is unexpected based on production by Ent-producing UTI89Δ*ybtS*Δ*iroA* (**Fig. 2B**). The nature of this discrepency is unclear and may arise from unappreciated catabolic differences, regulatory pathways, or intracellular trafficking connected to these different strains, the different cultures conditions, or combinations thereof.

**Figure 4.**
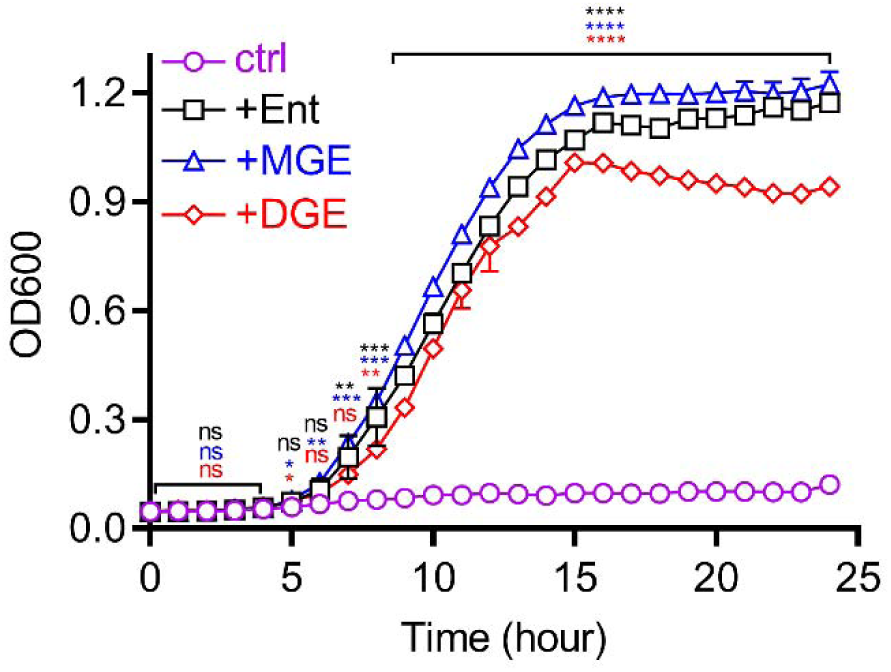
Trimer supplementation supports siderophore-null UTI89 mutant growth in siderophore-dependent growth medium. Growth of the siderophore-null strain UTI89Δ*entB*Δ*ybtS* was measured by optical density at 600 nm (OD600) in siderophore-dependent medium following supplementation with Ent, MGE, or DGE and compared to unsupplemented control (ctrl). Statistics were performed using 1-way ANOVA with Dunnett’s multiple-comparison test with P ≤ 0.05 considered as statistically significant. ns: not significant. *: *P* <= 0.05. **: *P* < 0.01. ***: *P* < 0.001. ****: *P* < 0.0001.

**Figure 5.**
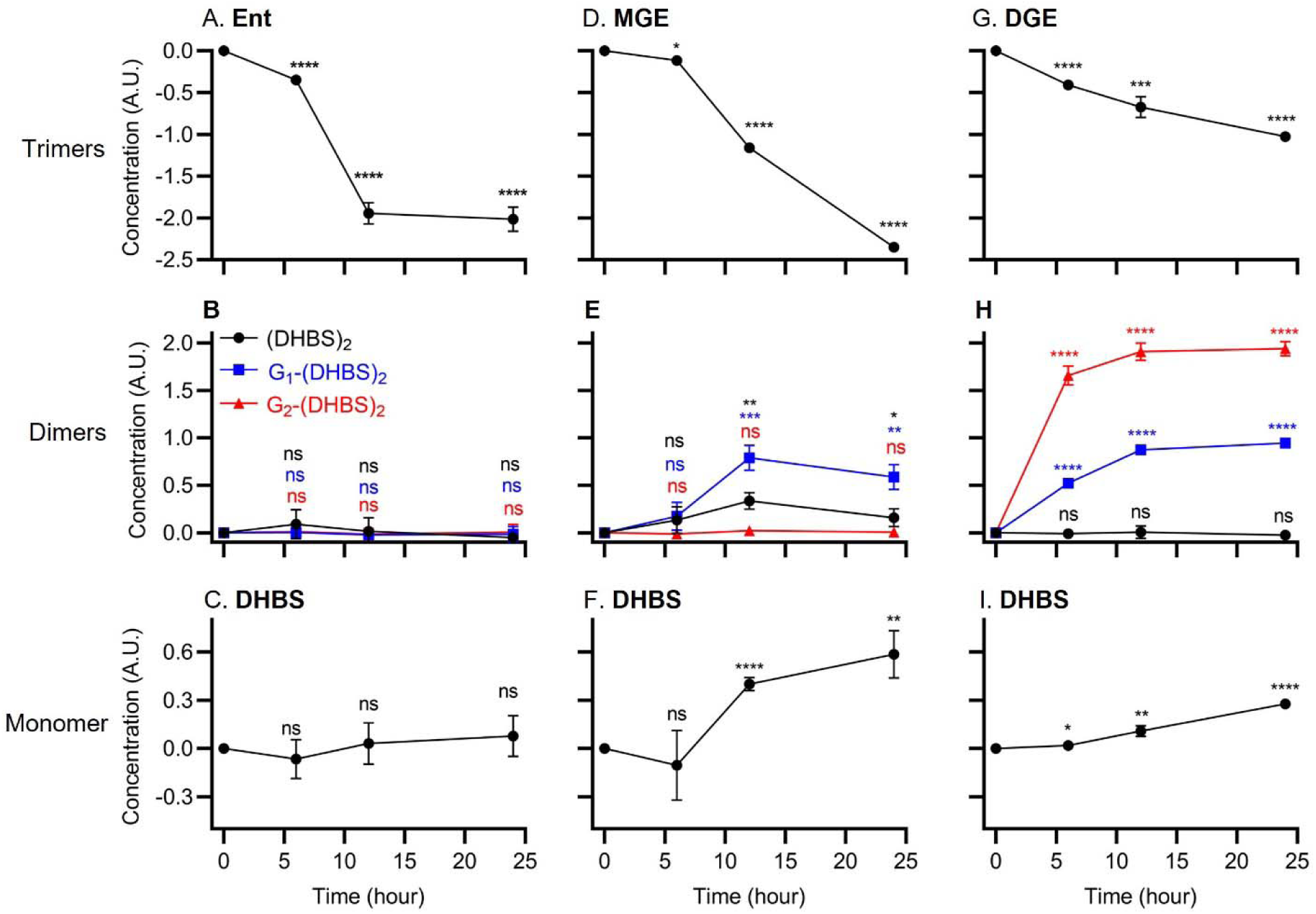
Enterobactin-associated exometabolites during trimer-dependent growth. Siderophore-null strain UTI89Δ*entB*Δ*ybtS* was cultured in siderophore-dependent medium containing purified Ent, MGE, or DGE. The enterobactin-associated metabolome in the medium was measured using LC-MS/MS at time points during culture. **(A & B & C)** Ent **(A)** is imported and catabolized by UTI89Δ*entB*Δ*ybtS* without producing any dimer **(B)** or monomer **(C)** *ent* catechol compounds. **(D & E & F)** MGS (D) is imported and catabolized by UTI89Δ*entB*Δ*ybtS*, which produces (DHBS)_2_ and G_1_-(DHBS)_2_ dimers **(E)** and DHBS monomer **(F)**. **(G & H & I)** DGS **(G)** is imported and catabolized by UTI89Δ*entB*Δ*ybtS*, which produces G_1_-(DHBS)_2_ and G_2_-(DHBS)_2_ dimers **(H)** and DHBS monomer **(I)**. Statistics were performed using unpaired *t* test with P ≤ 0.05 considered as statistically significant. ns: not significant. *: *P* <= 0.05. **: *P* < 0.01. ***: *P* < 0.001. ****: *P* < 0.0001.

### UTI89 uses exogenous dihydroxybenzoic acid (DHB) to synthesize enterobactin

It is unclear why UTI89 foregoes complete catabolic reclamation of intracellular trimer constituents to instead release incompletely hydrolyzed trimer catabolites to the extracellular space. Bonacorsi *et al.* have connected enhanced bacterial DHB production for siderophore biosynthesis as a virulence-associated activity in neonatal meningitis-associated *E. coli*, suggesting that UPEC could similarly benefit from DHB reclamation (59). To determine whether UPEC can use exogenously derived DHB to support trimer biosynthesis, we derived an experimental system to monitor its incorporation. Specifically, we used a reverse isotope-labelling strategy to detect incorporation of unlabeled carbon atoms from exogenous DHB during culture with ^13^C_3_-glycerol as the carbon source. We found that addition of 200 μM DHB led to the appearance of a new Ent isotopologue with a *m/z* value 21 atomic mass units lower than ^13^C-substituted enterobactin, consistent with ^12^C_7_-DHB incorporation at all three catechol sites (**Fig. 6, Table S2**). DHB supplementation also yielded lower levels of singly and doubly substituted isotopologues that are 7 and 14 atomic mass units lower, respectively (**supplemental Fig. S16, Table S2**). These results demonstrate the capacity for exogenous DHB incorporation into Ent biosynthesis by UTI89, demonstrating the potential value of complete Ent hydrolysis to UPEC.

**Figure 6.**
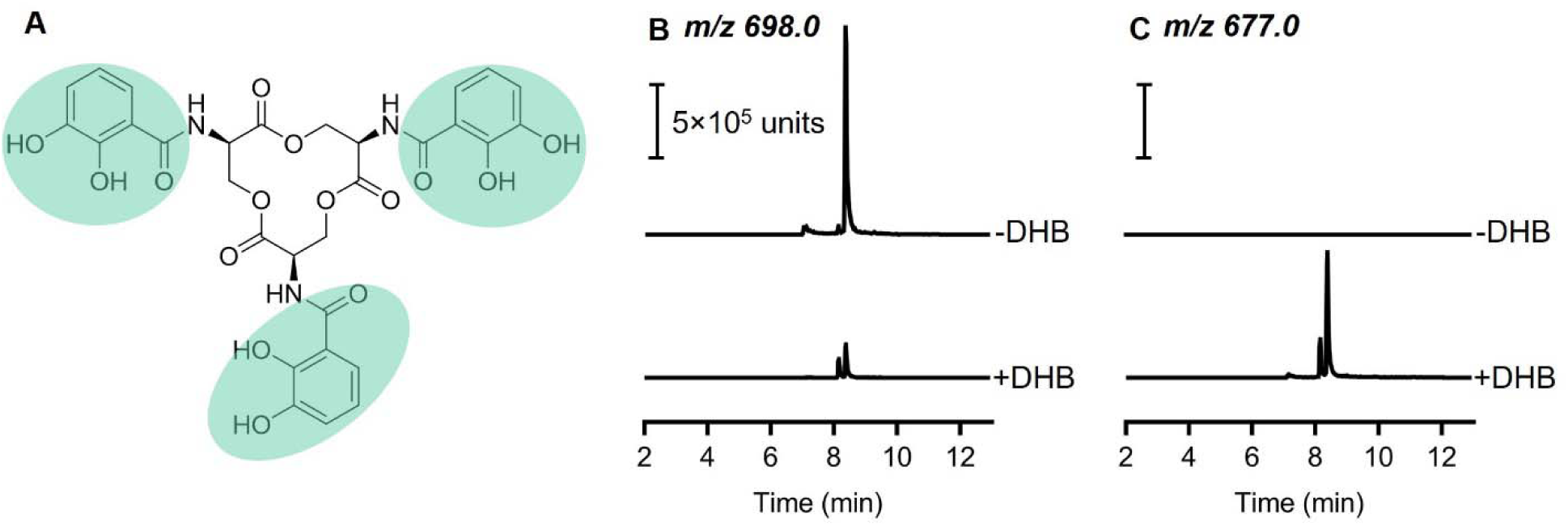
Exogenous 2,3-dihydrobenzoic acid (DHB) supports enterobactin biosynthesis. **(A)** Chemical structure of cyclic enterobactin (Ent) with the three DHB-derived groups, containing seven carbon atoms, highlighted in green. **(B**) LC-MS/MS detection of fully ^13^C_30_-substituted Ent ([M-H]^-^, *m/z* 698) in UTI89-conditioned ^13^C_3_-glycerol culture medium without (-DHB), or with (+DHB), 200 µM unlabeled DHB. (**C**) LC-MS/MS detection of ^13^C_9_-substituted Ent ([M-H]^-^, *m/z* 677) into which three DHB molecules have been incorporated without (-DHB), or with (+DHB), 200 µM unlabeled DHB.

### Siderophore activity of purified dimers

We hypothesized that UPEC forego complete trimer hydrolysis because the resulting dimers retain valuable siderophore activity. This would enable biosynthesis of one trimer molecule to support multiple rounds of iron import. To test this, we evaluated the siderophore activity of purified dimers in the siderophore-dependent growth condition described above. We observed that supplementation with either of two dimer metabolites, (DHBS)_2_ or G_2_-(DHBS)_2_, restored bacterial growth in iron-deficient conditions, with slower growth kinetics for G_2_-(DHBS)_2_ dimer than those observed for (DHBS)_2_ dimer and trimers (**Fig. 7A**, **Fig. 4**). DHBS production was generated from (DHBS)_2_ supplementation and catabolism only (**Fig. 7B-E**). Glucosylated *N*-(2,3-dihydroxybenzoyl)serine (G_1_-DHBS), that was expected to be generated from G_2_-(DHBS)_2_ hydrolysis, was poorly resolved in the LC-MS/MS conditions used here, likely because its high hydrophilicity renders it poorly resolved in reversed phase liquid chromatography. Together, these data are consistent with siderophore activity by both C-glucosylated and non-glycosylated dimers.

**Figure 7.**
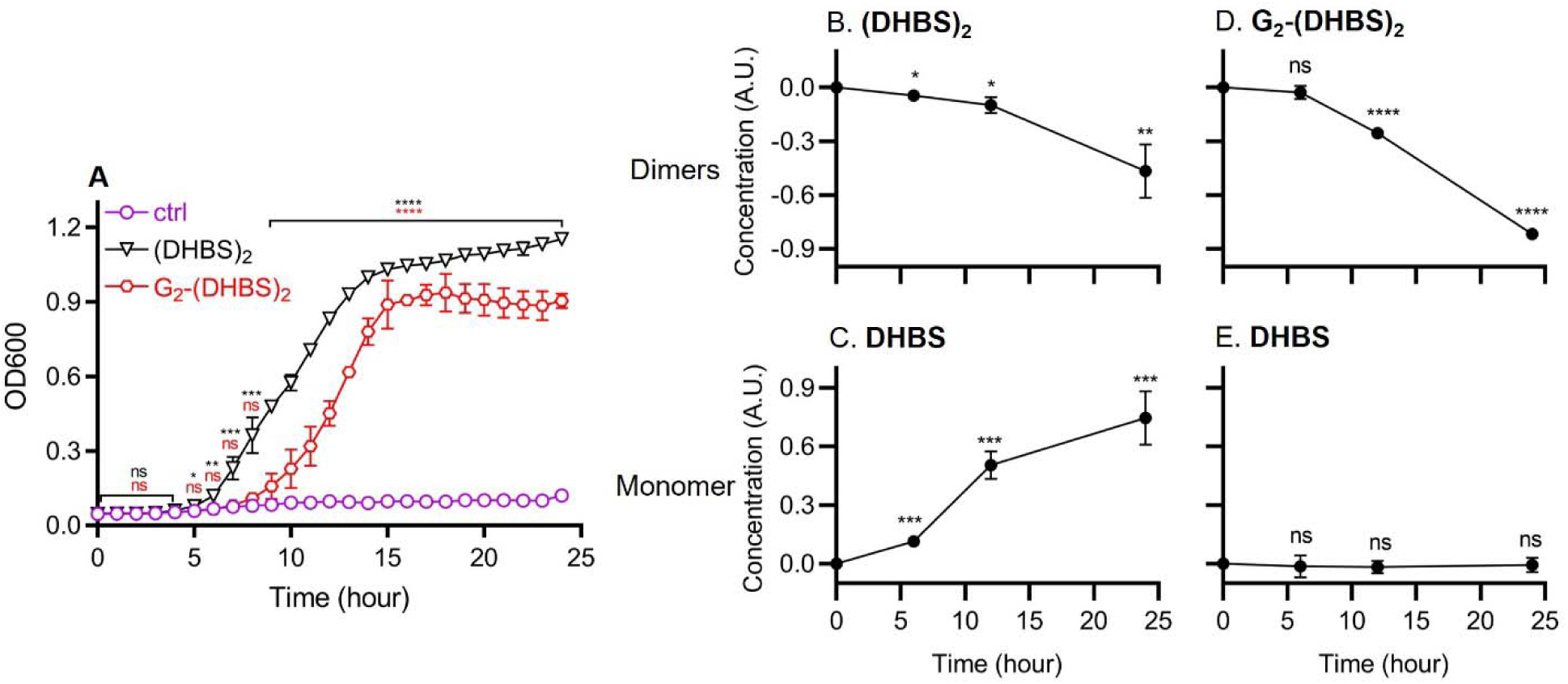
Enterobactin-associated dimers support siderophore-dependent growth. **(A)** Growth of the siderophore-null strain UTI89Δ*entB*Δ*ybtS* was measured by optical density at 600 nm (OD600) in siderophore-dependent medium following supplementation with the enterobactin-associated dimer exometabolites (DHBS)_2_ or G_2_-(DHBS)_2_, or siderophore-free control (ctrl). (**B-E**) The enterobactin-associated metabolome in the medium was measured using LC-MS/MS at time points during culture with dimer and monomer results shown for (DHBS)_2_-supplemented cultures (**B,C**) and G_2_-(DHBS)_2_.-supplemented cultures (**D,E**). Statistics were performed using unpaired *t* test with P ≤ 0.05 considered as statistically significant. ns: not significant. *: *P* <= 0.05. **: *P* < 0.01. ***: *P* < 0.001. ****: *P* < 0.0001.

## DISCUSSION

Multiple bacterial siderophore systems release exometabolites in addition to their canonical biosynthetic end products. Here, we find that UPEC have the potential to hydrolyze Ent trimers to recover raw materials for new biosynthesis, yet limit this process to instead generate and secrete incompletely hydrolyzed Ent (dimer), which is released as a siderophore. This suggests a bacterial “choice” between complete hydrolysis to maximize catabolic reclamation of biosynthetic substrates and incomplete hydrolysis to generate a dimeric catabolite that acts as a siderophores with lower iron(III) affinity. The former lowers the biosynthetic cost of new trimer biosynthesis while the latter yields another siderophore. The balance between these fates (complete or partial hydrolysis) may reflect evolutionary adaptation or, possibly, active regulation.

Siderophore function, as classically understood, is a metabolically costly process in which siderophore biosynthesis and secretion occurs because there is a chance some of these siderophores will diffuse back as iron complexes to support nutritional demands. For Ent and related siderophores, iron release requires hydrolysis by intracellular esterases, suggesting a diminished return on biosynthetic investment compared to siderophores that are non-destructively “recycled” and re-secreted (37,47). The results above suggest that a more nuanced situation has evolved in which trimer hydrolysis proceeds only to the extent necessary for iron release so that a catabolite may be secreted for additional rounds of siderophore-mediated iron delivery. The growth-promoting siderophore activity of dimers observed here is supported by a previous report of (DHBS)_2_-mediated ^55^Fe localization to *E. coli* (60). Additional supportive evidence was reported for iron-dependent growth of *Campylobacter jejuni*, an Ent non-producer that uses (DHBS)_2_ from *E. coli* as a siderophore in an example of siderophore “piracy” by this organism (61). The siderophore activity of dimers is thus associated with another example of metabolic cost-avoidance.

Enterobactin-associated dimers may have lower iron affinity than trimers, given the role of catechols in iron (III) affinity. A 1:1 dimer-iron complex provides catechol hydroxyl ligands for four of the six iron (III) coordination sites. This tetra-dentate coordination is observed for other siderophores such as pyochelin from *Pseudomonas aeruginosa* (62,63) and azotochelin from *Azotobacter vinelandii* (64). Despite this potential drop in affinity, we observed comparable iron acquisition capability by (DHBS)_2_ dimers and trimers, though differences could emerge in more stringent competitive iron binding conditions. As with DGE, G_2_-(DHBS)_2_ may similarly resist sequestration by Lipocalin-2 (14,33,34), rendering it more capable of delivering iron to UPEC during infections.

Incomplete trimer hydrolysis by UPEC suggests that siderophore esterase and peptidase systems have evolved to be inefficient as a means to support dimer-associated iron acquisition. The extent of hydrolysis may vary with the specific trimer and the hydrolases recruited during iron recovery. We found that siderophore-null UTI89 consumed purified Ent without significant monomer or dimer secretion, while purified glucosylated Ent trimers (MGE and DGE) resulted in abundant monomer and dimer formation. It remains unclear whether these differences reflect higher order metabolic interactions or unappreciated regulatory process affecting hydrolase activity, possibly responsive to Ent C-glucosylation. Complete hydrolysis of C-glucosylated trimers could be evolutionarily disfavored due to the likely inability to use C-glucosylated DHB as an Ent biosynthetic pathway. Lin, *et al.* previously reported that purified Fes and IroD can hydrolyze Ent trimer to produce DHBS monomer and (DHBS)_2_ dimer, and IroE can hydrolyze Ent to produce (DHBS)_2_ dimer (47).

We found that generation of short-length enterobactin-associated metabolites from trimers was dependent on TonB-dependent transport, consistent with their catabolic origin. It is likely that dimers similarly require TonB-dependent transporters, which may include transporters different from the Ent or glucosylated Ent transporters (Fep and IroN, respectively) associated with trimer transport. Fiu and Cir have been proposed to import Ent metabolites and catechols (65–68), with Cir serving as an important drug target for the catechol-modified cephalosporin cefiderocol (69–71), consistent with its activity in human infections. The substrate specificity for these transporters and their relationship to the network of enterobactin-associated exometabolites described here is incompletely understood and may yield deeper functional insights into enterobactin system function.

In conclusion, the exometabolite network described here is consistent with a series of regulatory and functional adaptations that minimize costs of Ent-mediated iron delivery in *E. coli* cells. Enterobactin biosynthesis, a metabolically costly process, is activated under iron-restricted conditions by Fur repressor regulation. At low bacterial density, *E. coli* have the ability to render Ent a private good, available only to the producing organism, and minimizing diffusional loss (72). Sub-maximal siderophore hydrolysis in UPEC to release dimers extends the iron delivery potential of Ent and its derivatives. Together, these activities results are consistent with a biochemical network connecting intracellular and extracellular *E. coli* metabolomes to cost-effectively support iron-dependent growth. These findings may help explain why enterobactin expression can be sustained as the universal siderophore system in urinary *E. coli* isolates. Aspects of this network may be useful in devising new antimicrobial therapeutics for uropathogenic *E. coli* and related bacteria.

## METHODS AND MATERIALS

### Bacterial strains and culture conditions

We examined exometabolite production, consumption, and use with the well-characterized, cystitis-derived model UPEC strain UTI89 and its previously described isogenic mutants UTI89Δ*entB*, UTI89Δ*tonB*, and UTI89Δ*entB*Δ*ybtS* (**Table S3**) (11,13,21,73). UPEC strain CFT073 was used for bacterial secondary metabolite production due to its high yield of C-glucosylated products (**Table S3**) (11, 13). Bacterial cultures were grown from single colonies in LB broth for overnight under 37□, washed with PBS, back-diluted 1:1000 into filter-sterilized M63 minimum media, inoculated with 200 µL into 96-well microplates, and incubated under 37□ for in the indicated assays. Experimental cultures were conducted in M63 minimum media containing 0.2% glycerol as a carbon source and 10 µg/mL nicotinic acid (low iron), with 100 µM FeCl_3_ (high iron), or with 10 µM bovine serum albumin addition (siderophore-dependent)(21, 32). Bacterial growth was quantified by the optical density at 600 nm (OD600) using a Spectrophotometer (Beckman Coulter, DU-800) or an incubated microplate reader (Tecan Spark).

### Untargeted liquid chromatography mass spectrometry (LC-MS)

Untargeted full scan LC-MS profiling was performed to characterize the extracellular metabolome (exometabolome) in media conditioned by UTI89 and UTI89Δ*entB* under low and high iron conditions. Conditioned medium was collected by centrifugation and filtration through 0.22 µM filters with storage at -80□. Samples were thawed on ice for liquid chromatography mass spectrometry (LC-MS) analysis with a Shimadzu Prominence UFLC-coupled AB Sciex 4000 QTrap mass spectrometer with a Turbo V electrospray ionization (ESI) source. LC separation was performed on an Ascentis Express phenyl-hexyl column (100 × 2.1 mm, 2.7 μm; Sigma-Aldrich) with solvent A (HPLC-grade water + 0.1% Formic acid; Sigma-Aldrich, Fluka) and B (90% Acetonitrile + 0.1% Formic acid; Sigma-Aldrich, Fluka) at 0.35 ml/min in a 36 minute gradient as follows: solvent B increased from 2% to 35% by 23 min, then increased to 98% by 33 min, and finally held steady at 98% for another 3 min. ESI-MS was performed in negative ion-enhanced MS mode, scanning from 50 to 1,500 *m/z*. A quality control sample was injected first and every ten samples thereafter to assess instrument stability. MarkerView, version 1.2.0 (Sciex) was used for peak alignment, generating the list of peaks for computational metabolome comparison analysis in the next section (13,14,32).

### Computational metabolomic comparison

Exometabolome comparisons between four groups of samples, including UTI89 grown in low and high iron media (wild type and wild type+Fe, respectively) and the enterobactin-null mutant UTI89Δ*entB* in low and high iron media (*entB* and *entB* +Fe, respectively), was performed on a combined computational model consisting of a sparse principal component analysis (sPCA) followed by a logistic regression (LR) classification. The computation was performed in R and Python, using the scikit-learn module and mixOmics package, respectively (74–77). Of note, sparsity penalization was enforced in the PCA dimensionality reduction step to prevent overfitting for this metabolome metadata consisting of much higher component dimensions than the number of samples (78,79). The iron-responsive sub-metabolome in UTI89 extracellular space were identified by the loadings analysis of all identified metabolites.

### Product ion scan and targeted LC-MS/MS

Product ion scan measurements were conducted to characterize chemical structures of the 10 enterobactin-associated molecules. The LC separation as above but with a flow of 0.5 ml/min and a 16 minute gradient as follows. Solvent B increased from 5% to 56% by 10 min, then increased to 98% by 12 min, and finally held steady at 98% for another 4 min. MS/MS product ion spectra of each negative ion was obtained in the enhanced product ion mode (80,81). Targeted LC-MS/MS multiplexed selected reaction monitoring (MRM) analyses were performed to validate the identities of 10 enterobactin-associated metabolites that were determined by the full-scan comparative metabolomic analysis as described above. MRM parameters protocols (**Table 1**) were established based on the results of product ion scan for each of the 10 targeted enterobactin-associated metabolites (11,13,14,21).

### Exometabolite purifications

Enterobactin-associated exometabolites were generated by growing CFT73 in M63/0.2% glycerol medium supplemented with 2,3-dihydroxybenzoic acid (DHB; Sigma-Aldrich) and 100 µM Dipyridyl at 37□ for 18 hours. Culture supernatant was collected and separated by separated by four consecutive steps, including a DEAE-sepharose resin (Sigma), an Amberlite XAD16N resin (20 ∼ 60 mesh, Sigma), an Kromasil Eternity 5-PhenylHexyl column (250 × 4.6 mm, 5 μm; Nouryon), and an Ascentis Express Phenyl-Hexyl column (100 × 4.6 mm, 2.7 μm; Sigma-Aldrich) to achieve the purification of five enterobactin-associated molecules, including Ent, MGE, DGE, [(DHBS)_2_, and G_2_-(DHBS)_2_, as previously described (11,47). Culture supernatant was first applied to a methanol (20%)-conditioned DEAE-sepharose column (Sigma). The column was washed with water and then eluted with 7.5 M ammonium formate. The DEAE eluate was supplemented with 120 mM sodium dithionite, incubated with methanol-conditioned Amberlite resin (XAD16N, Sigma-Aldrich) overnight, and eluted with 100% methanol. The eluate was concentrated in a rotatory evaporator (R-100 Rotavapor, BUCHI), lyophilized (Labconco), resuspended in HPLC-grade water plus 0.1% formic acid, and further purified on a Bio-Rad BioLogic DuoFlow 10 system equipped with a QuadTec UV-Vis detector and a Kromasil Eternity-5-PhenylHexyl column (Sigma-Aldrich). The Kromasil column was run at 0.30 ml/min with HPLC-grade water plus 0.1% formic acid (solvent A) and acetonitrile plus 0.1% formic acid (solvent B) using gradient as follows. Solvent B held steady at 2% for 1.0 ml, then increased to 15% over 1 ml, then increased to 52% over 40 ml, and finally increased to 100% over 1 ml. The DuoFlow elute was finally separated by another Ascentis Express Phenyl-Hexyl column (Sigma-Aldrich) in a Shimadzu Prominence UFLC system coupled with an SPD-M20A Prominence Diode Array (PDA) Detector. In order to purify the compounds with different properties, the LC separation was performed by injecting solvent A (HPLC-grade water + 0.1% Formic acid; Sigma-Aldrich, Fluka) and B (90% Acetonitrile + 0.1% Formic acid; Sigma-Aldrich, Fluka) at 0.5 ml/min with an 44 minute gradient under two scenarios as follows. Solvent B increased from 2% to 35% or 44% by 35 min, then increased to 98% B by 38 min, and finally held steady at 98% for another 6 min. Fractions containing purified molecules were measured via UV-Vis detection at 319 nm, pooled together, dried down by lyophilization, and stored in -80 □ freezer. On day of use, samples were resuspended in HPLC-grade water plus 0.1% formic acid, and concentrations were calculated by Beer-Lambert law using UV-Vis absorbances at 319 nm with an extinction coefficient of 11200 M^-1^cm^-1^. Purity was confirmed by targeted LC-MS/MS measurements (11,50,83,84).

### Exogeneous DHB for synthesizing enterobactin in isotope-labelling assay

To determine whether UPEC can synthesize Ent from DHB that is not immediately generated by endogenous DHB biosynthesis (40,41,85), we grew UTI89 from single colonies in LB broth for 12 h at 37□ was washed with PBS, back-diluted 1:1000 into ^13^C_3_-glycerol M63 minimum media with or without the supplement of 200 uM ^12^C-DHB in a 96-well plate, and grown at 37 for 24 hours. Targeted LC-MS/MS of Ent was conducted to monitor the incorporation of ^12^C from incorporation of unlabeled DHB (**Table S2**) by comparing ^13^C-substituted enterobactin isotopologues into which 0, 1, 2, or 3 ^12^C_7_-DHB were incorporated.

### Data availability

The computer codes for the analyses in this study are available in Github (https://github.com/QL5001/EntMetabolome_script; branch name: main; commit ID, 82317f0). All other data generated and analyzed in this study are included in the published article and supplementary materials.

### Statistical methods

GraphPad Prism 9.0 (GraphPad software) was used to generate graphs and perform statistical analysis in this study. We used the unpaired, two-tailed *t* test for comparisons between two groups, and one-way ANOVA for multigroup comparisons. *P* < 0.05 was considered statistically significant.

## ACKNOWLEDGEMENTS

The authors acknowledge funding from the Centers for Disease Control Prevention Epicenters Program Grant (CU54 CK 000162). JPH acknowledges National Institutes of Health grants R01DK099534 and R01DK111930. The content is solely the responsibility of the authors and does not necessarily represent the official view of the CDC or NIH.

## AUTHOR CONTRIBUTIONS

ZZ and JPH originally developed the concept and designed the overall study approach. ZZ performed the experiments. ZZ and JIR designed the mass spectrometry approaches. ZZ and LKS designed and performed the HPLC purification of bacterial secondary metabolites. ZZ and JPH analyzed the data. ZZ and JPH wrote the manuscript.

