## Supporting information for "Uropathogenic *Escherichia coli* wield enterobactin-derived catabolites as siderophores"

**SUPPLEMENTARY FIGURES**

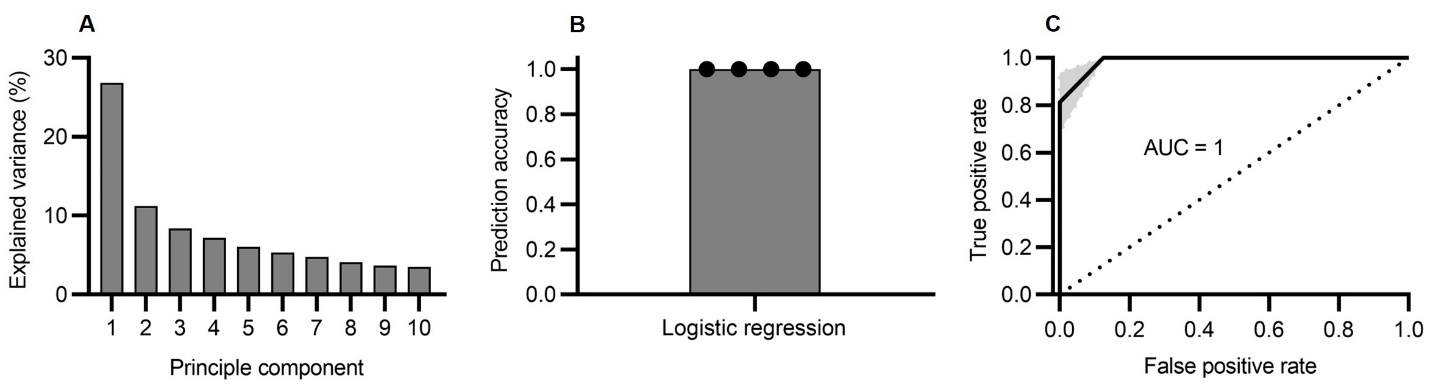

**Figure S1. Identification of enterobactin-associated compounds in uropathogenic *E. coli* (UPEC) UTI89 exometabolome. (A)** Explained variance of the first ten principal components from sparse principal component analysis (sPCA) of media conditioned by UTI89 grown in low and high iron media (wild type and wild type+Fe, respectively) and the enterobactin-null mutant UTI89∆*entB* in low and high iron media (*entB* and *entB* +Fe, respectively). High iron medium is achieved by addition of 100 µM FeCl_3_. **(B)** Logistic regression using sPCA-derived PC1 values for classifying between the group of “wild type” and the combination group of “wild type+Fe”, “*entB*”, and “*entB* +Fe” yielded a prediction accuracy of 1.0 (SD = 0) with 4-fold cross validations. **(C)** Logistic regression using sPCA-associated PC1 values for classifying between the group of “wild type” and the combination group of “wild type+Fe”, “*entB*”, and “*entB* +Fe” yielded an area under the curve (AUC) of 1.0 (SD = 0) with 4-fold cross validations.

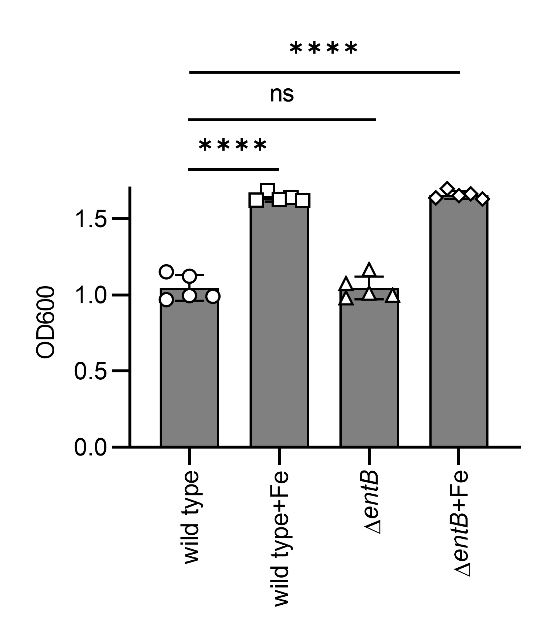

**Figure S2. Bacterial growths of wild type UTI89** in low and high iron media (wild type and wild type+Fe, respectively) and the enterobactin-null mutant UTI89∆*entB* in low and high iron media (*entB* and *entB* +Fe, respectively) after 18 hours incubation under 37°C were measured by optical density at 600 nm (OD600). Statistics were performed using 1-way ANOVA with Dunnett’s multiple-comparison test with P ≤ 0.05 considered as statistically significant. ns: not significant. *: *P* <= 0.05. **: *P* < 0.01. ***: *P* < 0.001. ****: *P* < 0.0001.

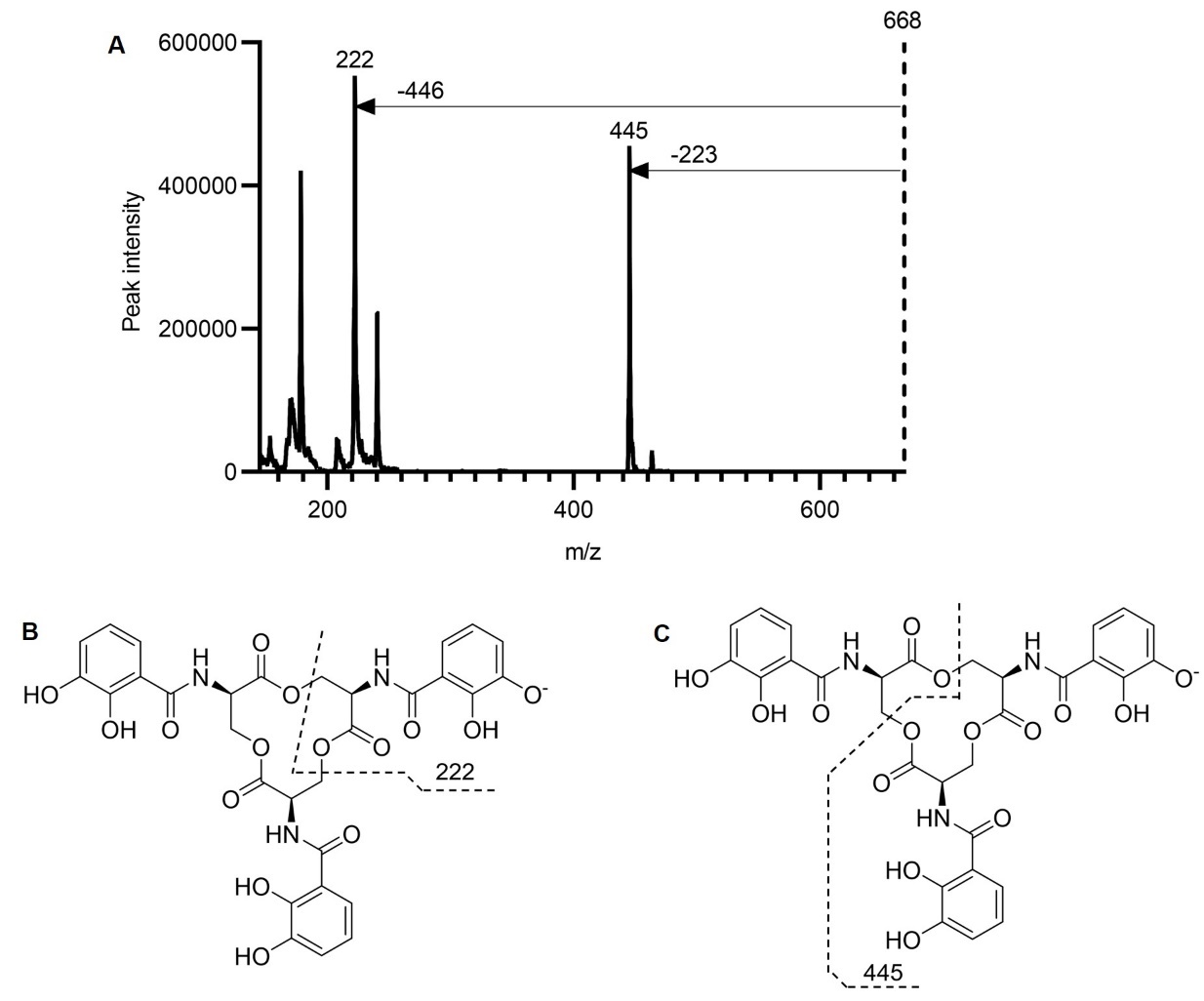

**Figure S3. Product ion scan mass spectrometry validates the chemical structure of enterobactin (Ent). (A)** Product ion spectrum of Ent. **(B)** The 222 Da fragment corresponded to a negative ion of (DHBS)^-^, with an empiric formula of C_10_H_8_NO_5_^-^. **(C)** The 445 Da fragment corresponded to a negative ion of (DHBS)_2_^-^, with an empiric formula of C_20_H_17_N_2_O_10_^-^.

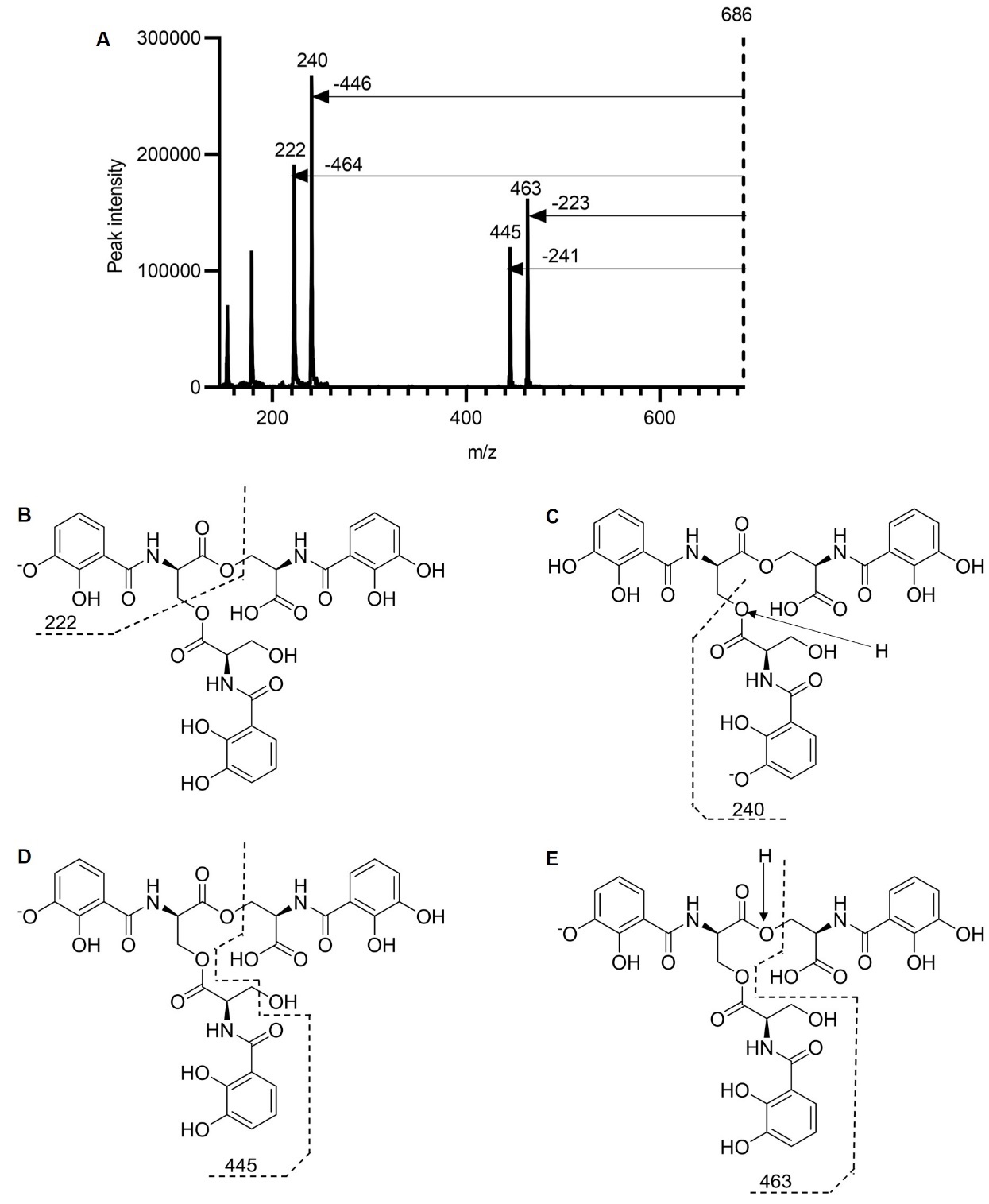

**Figure S4. Product ion scan mass spectrometry validates the chemical structure of linear enterobactin (lin-Ent). (A)** Product ion spectrum of lin-Ent. **(B)** The 222 Da fragment corresponded to a negative ion of (DHBS)^-^, with an empiric formula of C_10_H_8_NO_5_^-^. **(C)** The 240 Da fragment corresponded to a negative ion of (DHBS)^-^, with an empiric formula of C_10_H_10_NO_6_^-^. **(D)** The 445 Da fragment corresponded to a negative ion of (DHBS)_2_^-^, with an empiric formula of C_20_H_17_N_2_O_10_^-^. **(E)** The 463 Da fragment corresponded to a negative ion of (DHBS)_2_^-^, with an empiric formula of C_20_H_19_N_2_O_11_^-^.

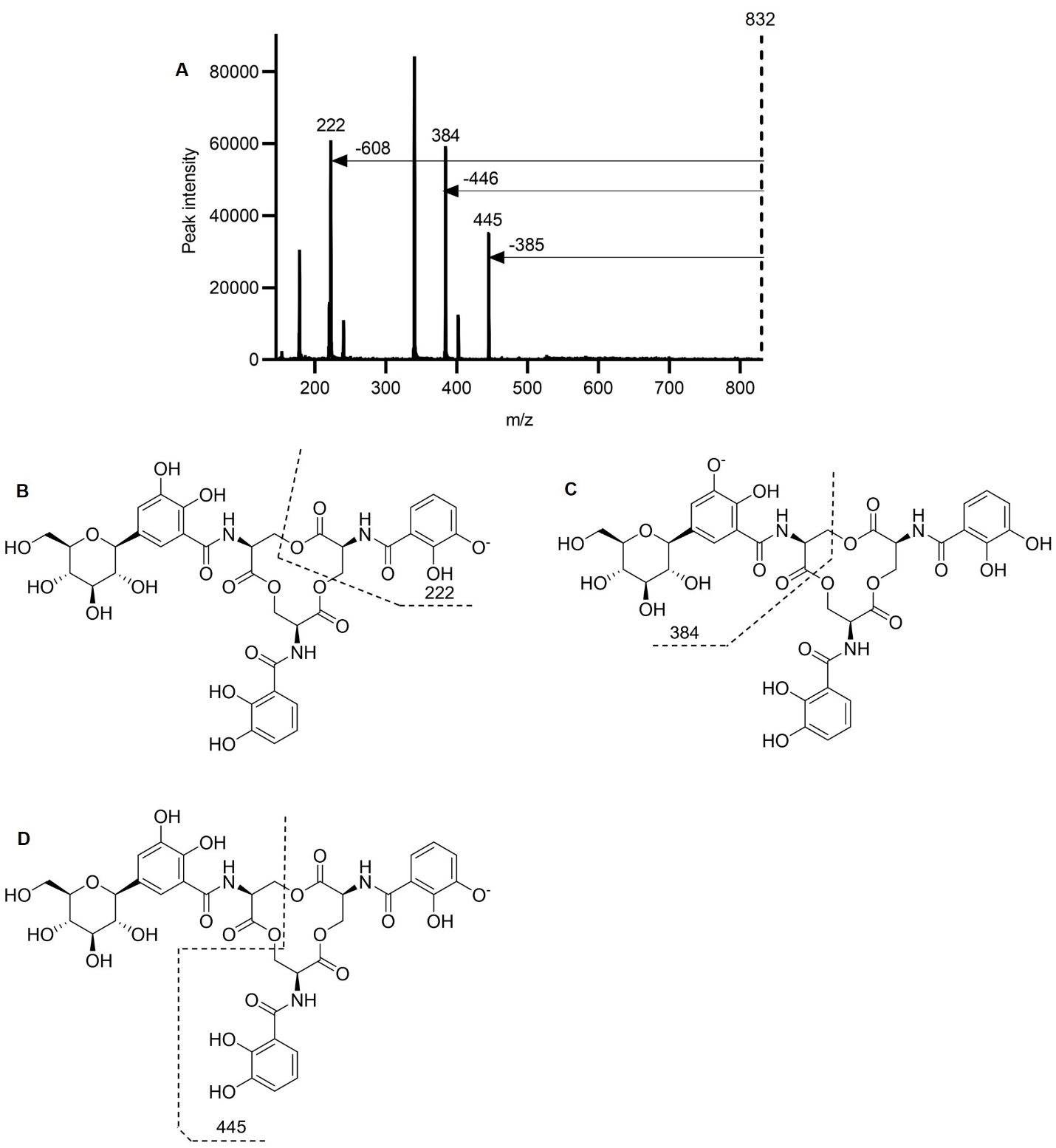

**Figure S5. Product ion scan mass spectrometry validates the chemical structure of monoglucosylated enterobactin (MGE). (A)** Product ion spectrum of MGE. **(B)** The 222 Da fragment corresponded to a negative ion of (DHBS)^-^, with an empiric formula of C_10_H_8_NO_5_^-^. **(C)** The 384 Da fragment corresponded to a negative ion of [G_1_-(DHBS)_1_]^-^, with an empiric formula of C_16_H_18_NO_10_^-^. **(D)** The 445 Da fragment corresponded to a negative ion of (DHBS)_2_^-^, with an empiric formula of C_20_H_17_N_2_O_10_^-^.

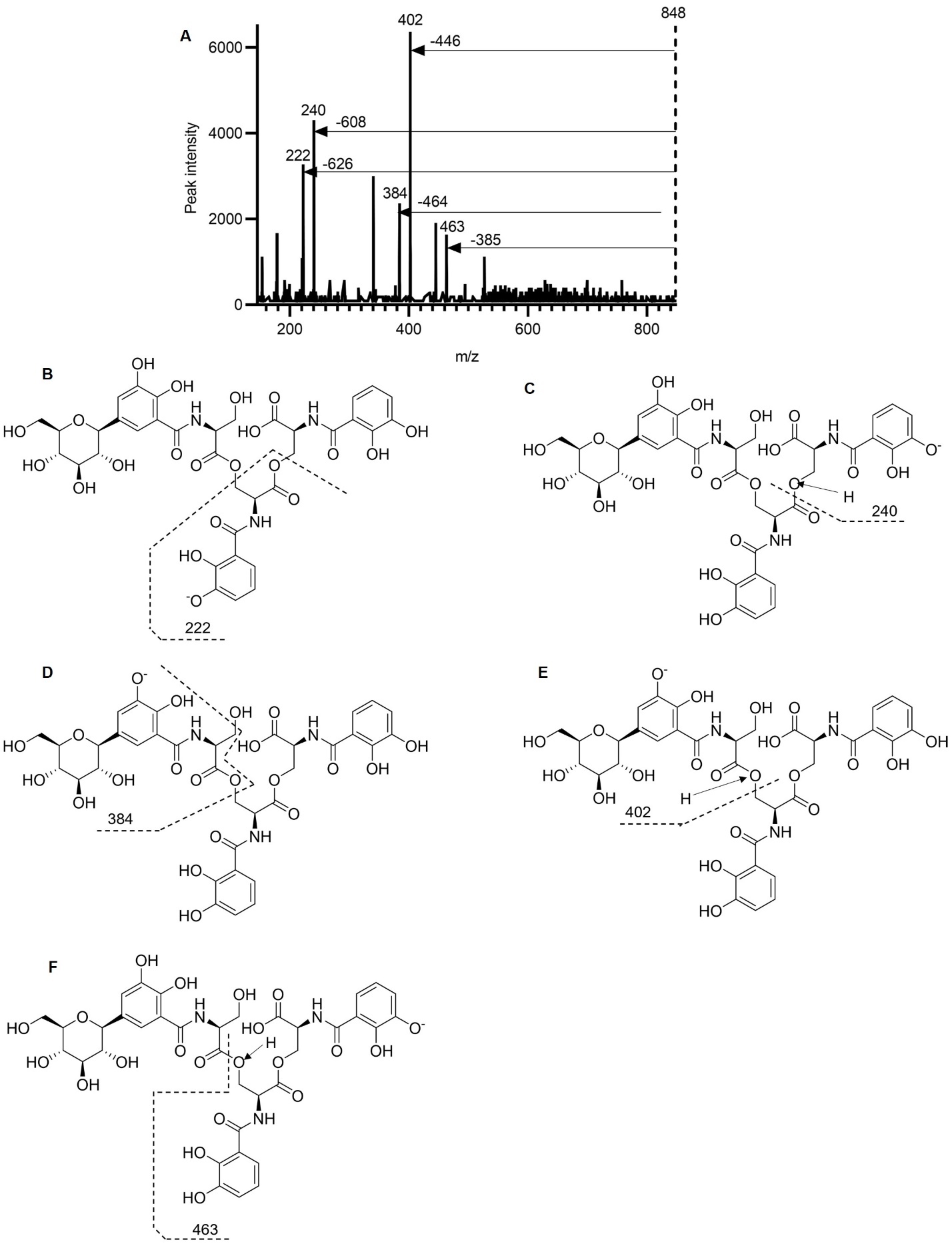

**Figure S6. Product ion scan mass spectrometry validates the chemical structure of linear monoglucosylated enterobactin (lin-MGE). (A)** Product ion spectrum of lin-MGE. **(B)** The 222 Da fragment corresponded to a negative ion of (DHBS)^-^, with an empiric formula of C_10_H_8_NO_5_^-^. **(C)** The 240 Da fragment corresponded to a negative ion of (DHBS)^-^, with an empiric formula of C_10_H_10_NO_6_^-^. **(D)** The 384 Da fragment corresponded to a negative ion of [G_1_-(DHBS)_1_]^-^, with an empiric formula of C_16_H_18_NO_10_^-^. **(E)** The 402 Da fragment corresponded to a negative ion of [G_1_-(DHBS)_1_]^-^, with an empiric formula of C_16_H_18_NO_11_^-^. **(F)** The 463 Da fragment corresponded to a negative ion of (DHBS)_2_^-^, with an empiric formula of C_20_H_19_N_2_O_11_^-^.

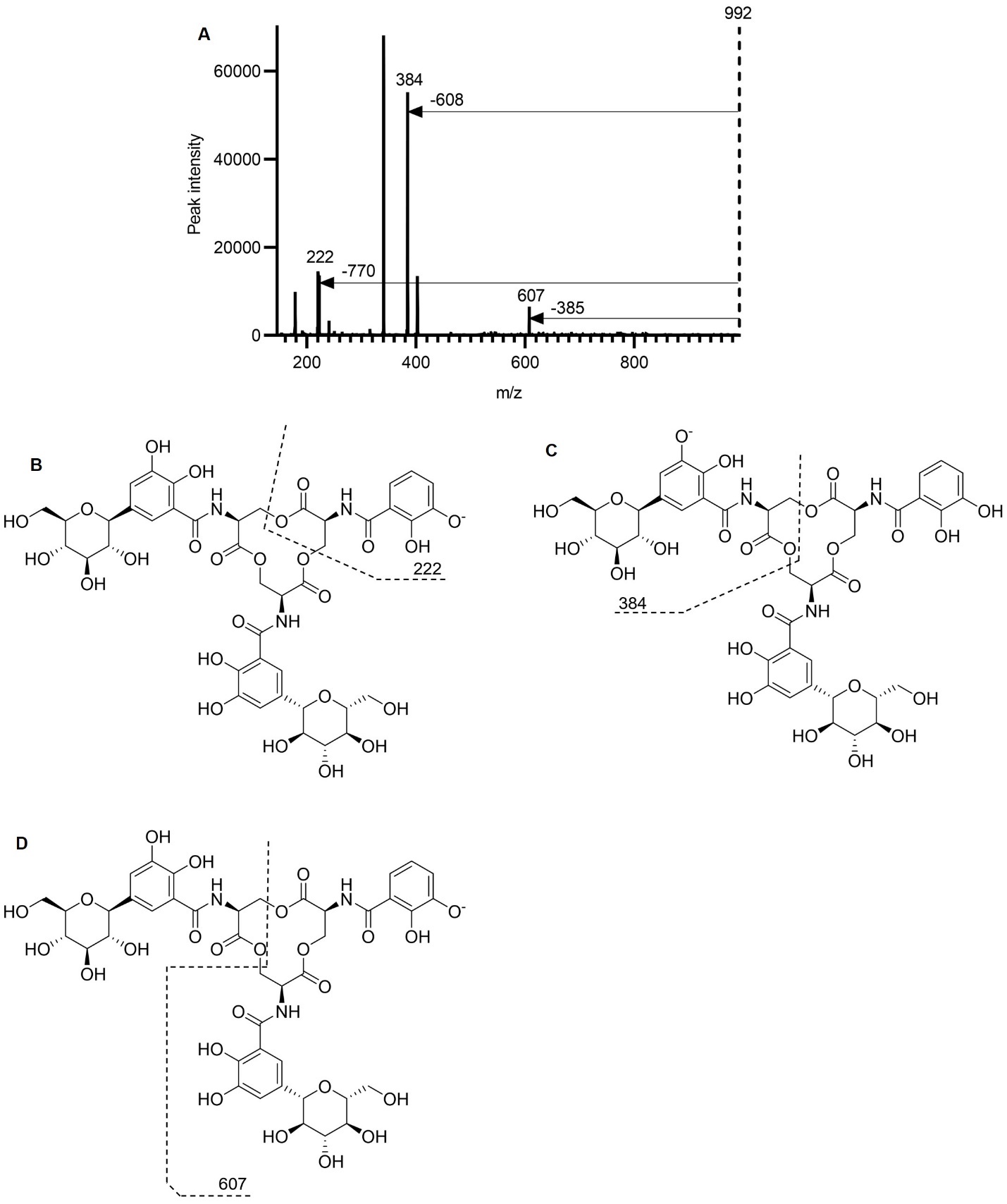

**Figure S7. Product ion scan mass spectrometry validates the chemical structure of diglucosylated enterobactin (DGE). (A)** Product ion spectrum of DGE. **(B)** The 222 Da fragment corresponded to a negative ion of (DHBS)^-^, with an empiric formula of C_10_H_8_NO_5_^-^. **(C)** The 384 Da fragment corresponded to a negative ion of [G_1_-(DHBS)_1_]^-^, with an empiric formula of C_16_H_18_NO_10_^-^. **(D)** The 607 Da fragment corresponded to a negative ion of [G_1_-(DHBS)_2_]-, with an empiric formula of C_26_H_27_N_2_O_15_^-^.

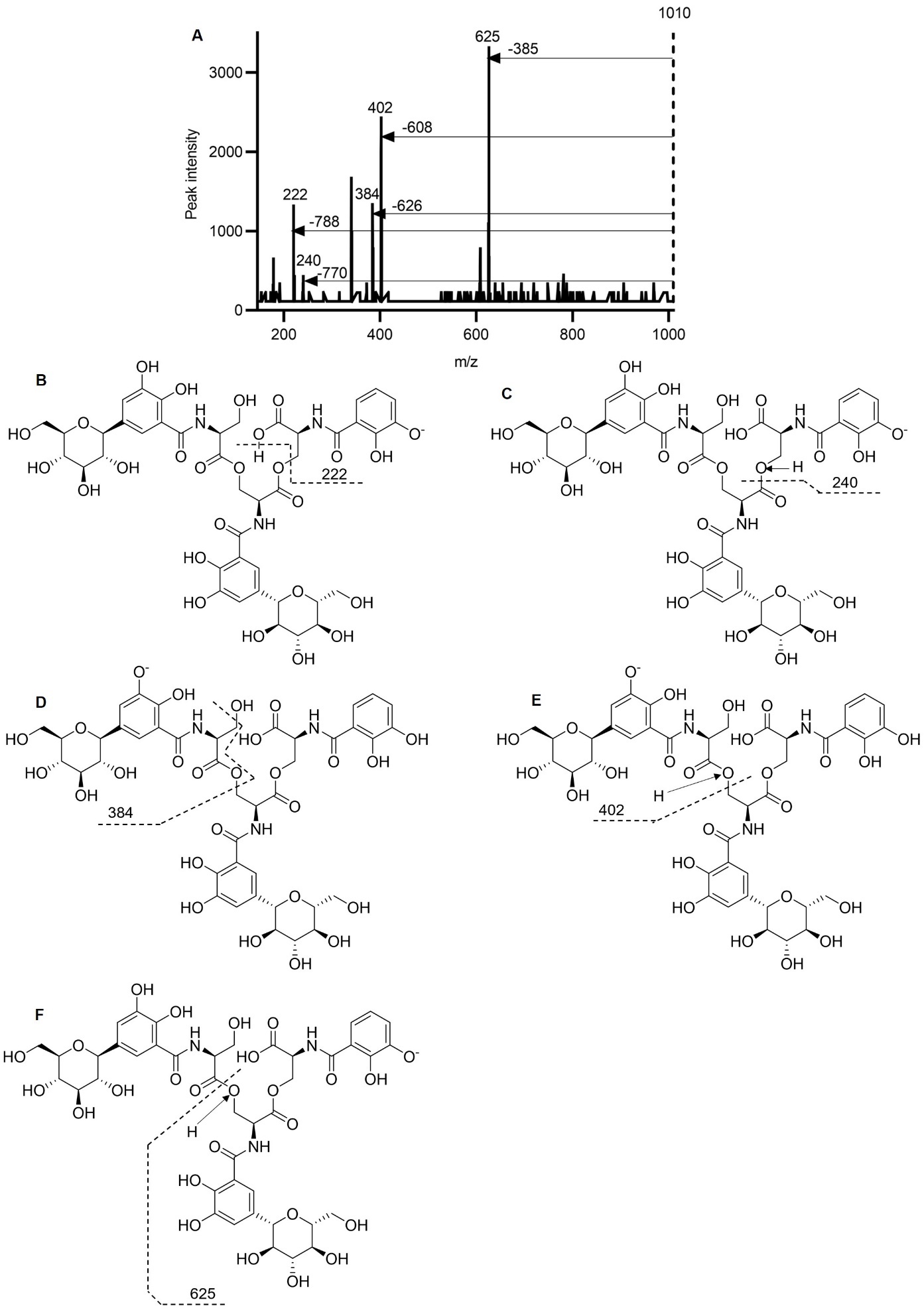

**Figure S8. Product ion scan mass spectrometry validates the chemical structure of linear diglucosylated enterobactin (lin-DGE). (A)** Product ion spectrum of lin-DGE. **(B)** The 222 Da fragment corresponded to a negative ion of (DHBS)^-^, with an empiric formula of C_10_H_8_NO_5_^-^. **(C)** The 240 Da fragment corresponded to a negative ion of (DHBS)^-^, with an empiric formula of C_10_H_10_NO_6_^-^. **(D)** The 384 Da fragment corresponded to a negative ion of [G_1_-(DHBS)_1_]^-^, with an empiric formula of C_16_H_18_NO_10_^-^. **(D)** The 607 Da fragment corresponded to a negative ion of [G_1_-(DHBS)_2_]-, with an empiric formula of C_26_H_27_N_2_O_15_^-^. **(E)** The 402 Da fragment corresponded to a negative ion of [G_1_-(DHBS)_1_]^-^, with an empiric formula of C_16_H_18_NO_11_^-^. **(F)** The 625 Da fragment corresponded to a negative ion of [G_1_-(DHBS)_2_]^-^, with an empiric formula of C_26_H_29_N2O_16_^-^.

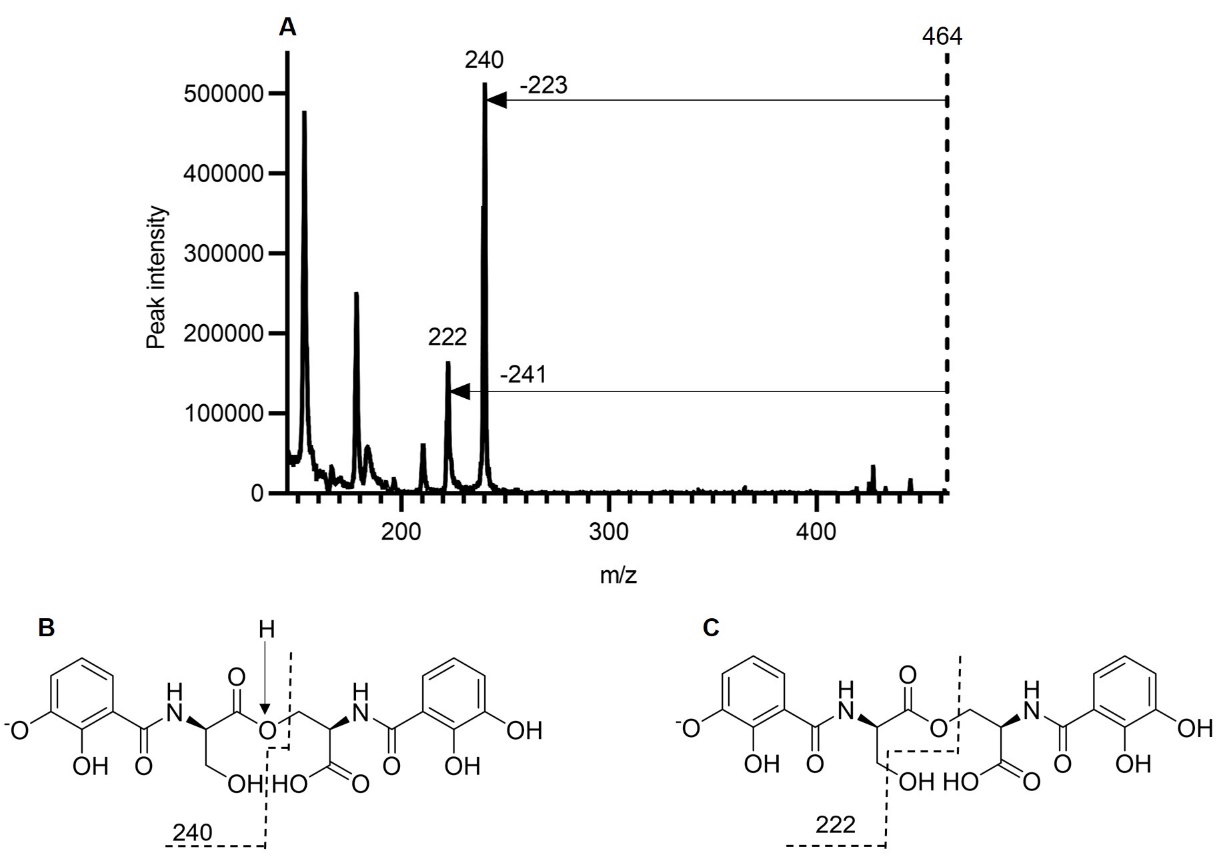

**Figure S9. Product ion scan mass spectrometry validates the chemical structure of (DHBS)_2_. (A)** Product ion spectrum of (DHBS)_2_. **(B)** The 222 Da fragment corresponded to a negative ion of (DHBS)^-^, with an empiric formula of C_10_H_8_NO_5_^-^. **(C)** The 240 Da fragment corresponded to a negative ion of (DHBS)^-^, with an empiric formula of C_10_H_10_NO_6_^-^.

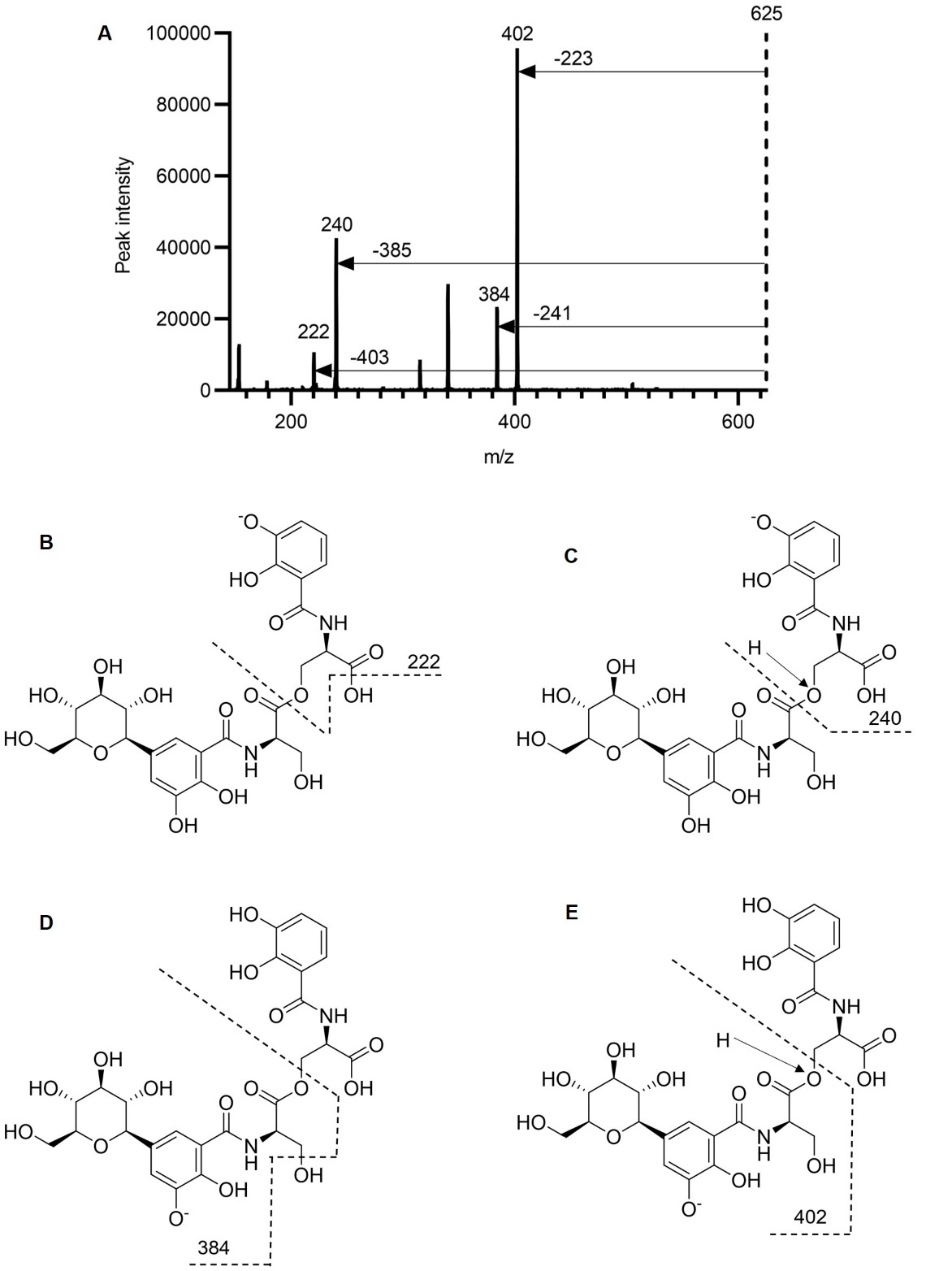

**Figure S10. Product ion scan mass spectrometry validates the chemical structure of G_1_-(DHBS)_2_. (A)** Product ion spectrum of G_1_-(DHBS)_2_. **(B)** The 222 Da fragment corresponded to a negative ion of (DHBS)^-^, with an empiric formula of C_10_H_8_NO_5_^-^. **(C)** The 240 Da fragment corresponded to a negative ion of (DHBS)^-^, with an empiric formula of C_10_H_10_NO_6_^-^. **(D)** The 384 Da fragment corresponded to a negative ion of [G_1_-(DHBS)_1_]^-^, with an empiric formula of C_16_H_18_NO_10_^-^. **(E)** The 402 Da fragment corresponded to a negative ion of [G_1_-(DHBS)_1_]^-^, with an empiric formula of C_16_H_18_NO_11_^-^.

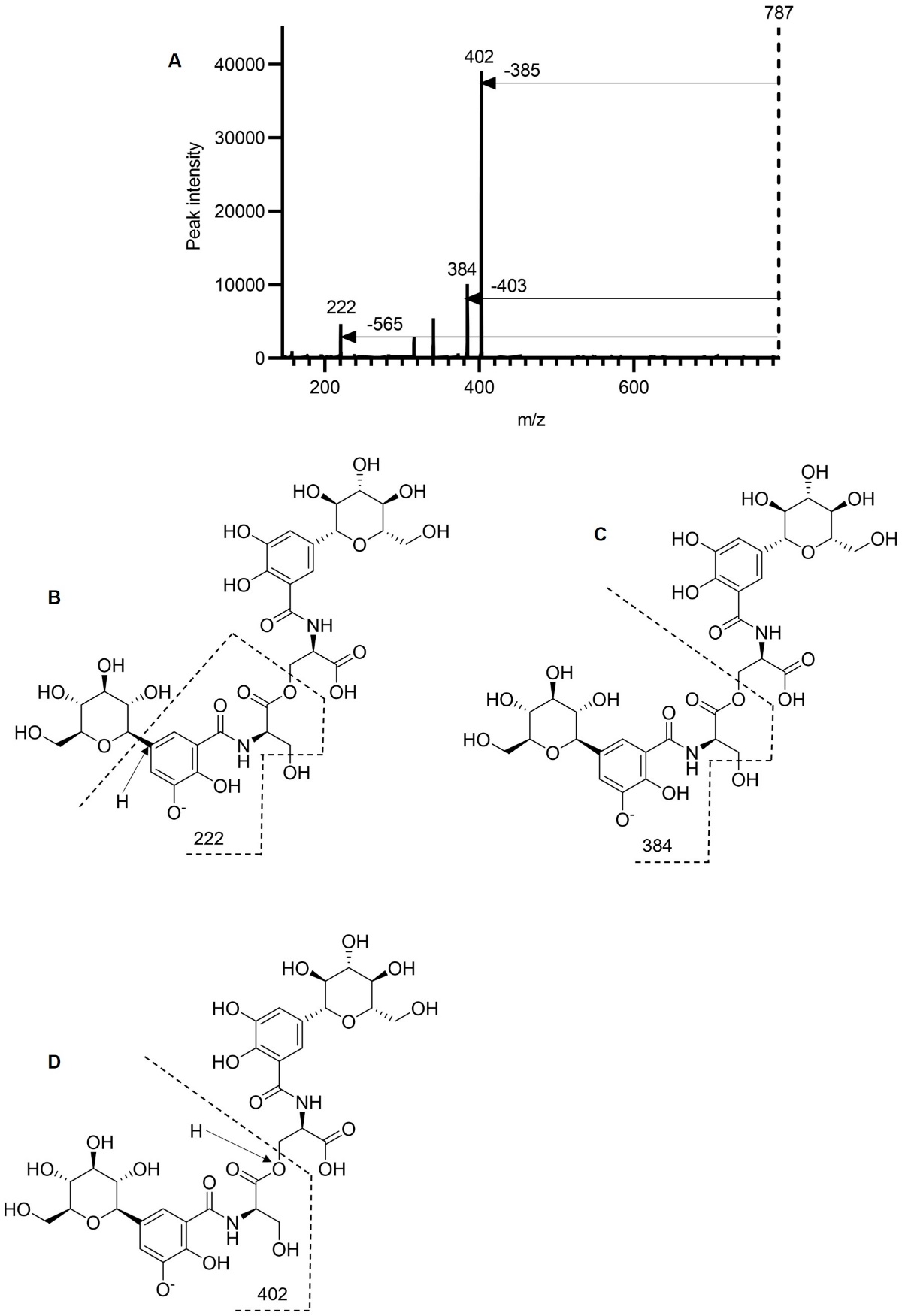

**Figure S11. Product ion scan mass spectrometry validates the chemical structure of G_1_-(DHBS)_2_. (A)** Product ion spectrum of G_2_-(DHBS)_2_. **(B)** The 222 Da fragment corresponded to a negative ion of (DHBS)^-^, with an empiric formula of C_10_H_8_NO_5_^-^. **(C)** The 384 Da fragment corresponded to a negative ion of [G_1_-(DHBS)_1_]^-^, with an empiric formula of C_16_H_18_NO_10_^-^. **(D)** The 402 Da fragment corresponded to a negative ion of [G_1_-(DHBS)_1_]^-^, with an empiric formula of C_16_H_18_NO_11_^-^.

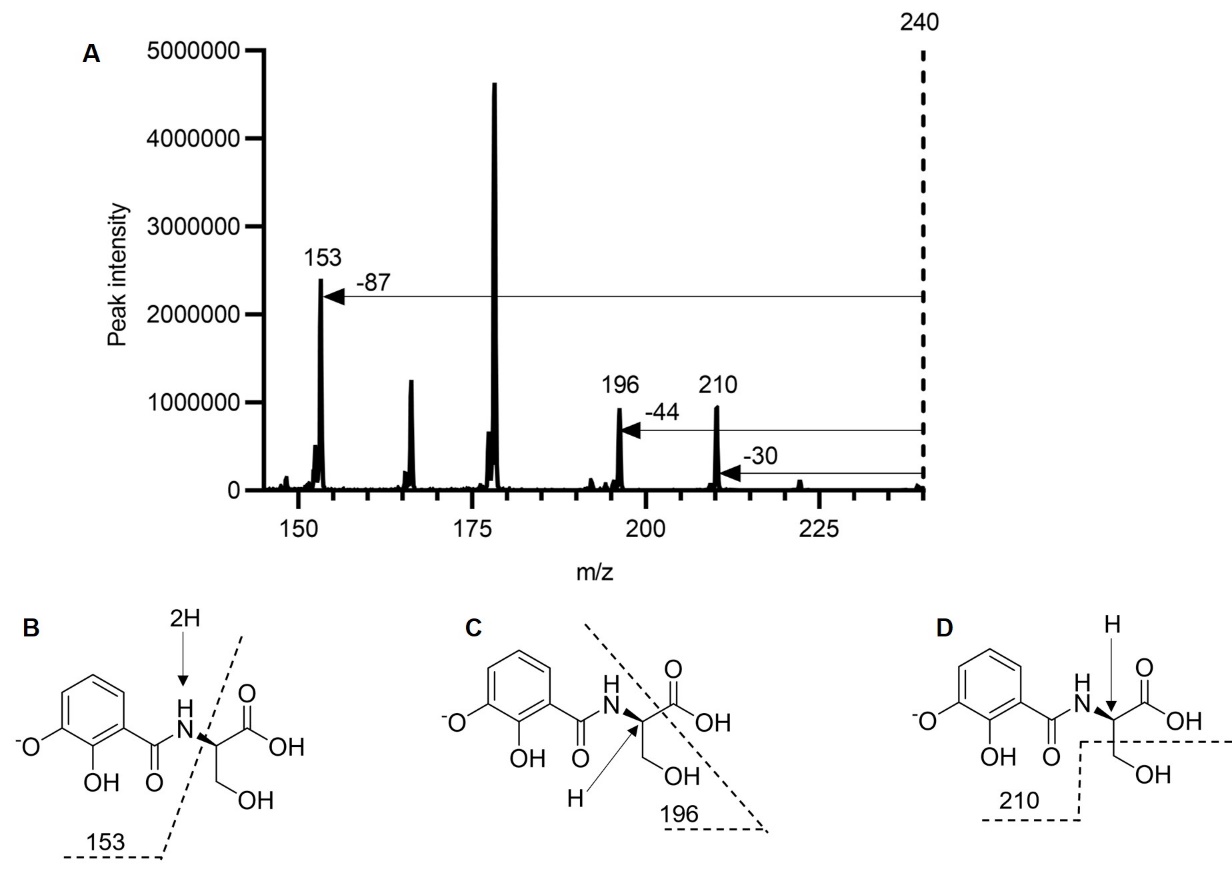

**Figure S12. Product ion scan mass spectrometry validates the chemical structure of DHBS. (A)** Product ion spectrum of DHBS. **(B)** The 153 Da fragment corresponded to a fragmented negative ion of (DHBS)^--^, with an empiric formula of C_5_H_7_NO_3_^-^. **(C)** The 196 Da fragment corresponded to a fragmented negative ion of (DHBS)^-^, with an empiric formula of C_9_H_10_NO_4_^-^. **(C)** The 210 Da fragment corresponded to a fragmented negative ion of (DHBS)^-^, with an empiric formula of C_9_H_8_NO_5_^-^.

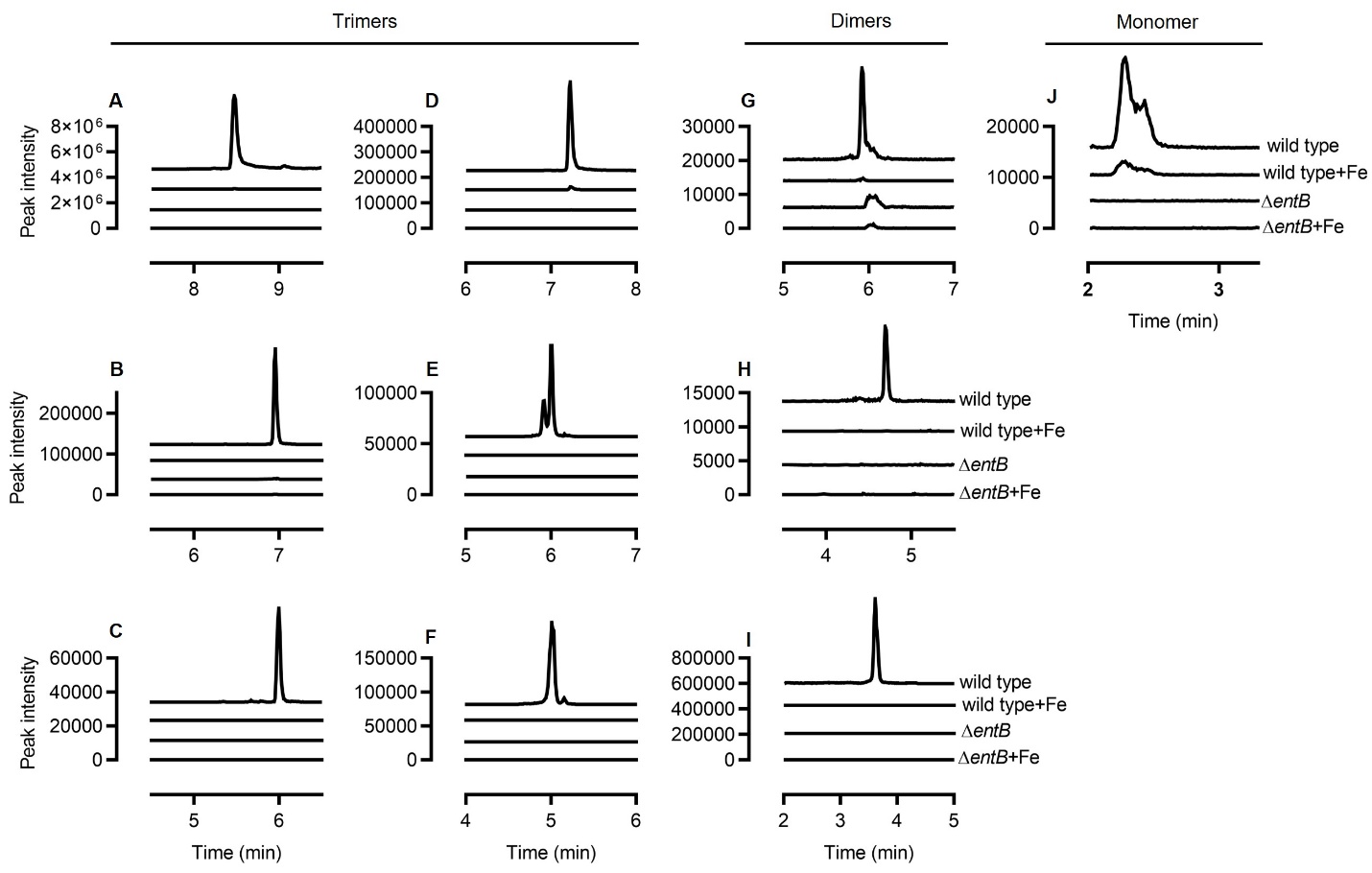

**Figure S13. Targeted LC-MS/MS detects a series of 10 enterobactin-associated *N*-(2,3-dihydroxybenzoyl)serine (DHBS) polymer compounds.** LC-MS/MS chromatograms corresponding to the precursor-product ions from **(A)** Ent, **(B)** lin-Ent, **(C)** MGE, **(D**) lin-MGE, **(E)** DGE, **(F)** lin-DGE, **(G)** (DHBS)_2_, **(H)** G_1_-(DHBS)_2_, **(I)** G_2_-(DHBS)_2_, and **(J)** DHBS for four experimental groups, including UTI89 grown in low and high iron media (wild type and wild type+Fe, respectively) and the enterobactin-null mutant UTI89∆*entB* in low and high iron media (*entB* and *entB* +Fe, respectively).

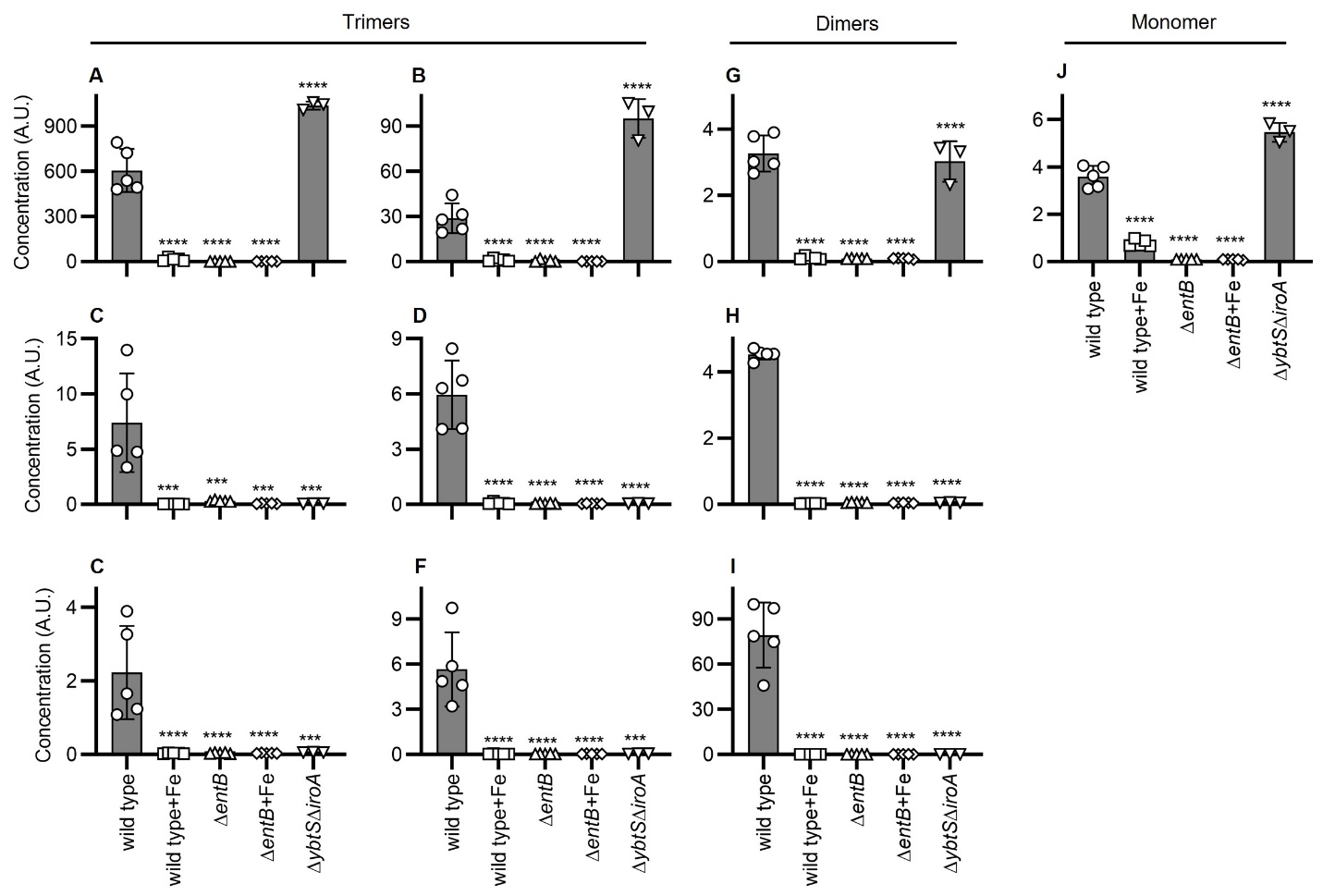

**Figure S14. Quantitative comparison for a series of 10 enterobactin-associated *N*-(2,3-dihydroxybenzoyl)serine (DHBS) polymer compounds using targeted LC-MS/MS.** LC-MS/MS quantification and comparison for the precursor-product ions from **(A)** Ent, **(B)** lin-Ent, **(C)** MGE, **(D**) lin-MGE, **(E)** DGE, **(F)** lin-DGE, **(G)** (DHBS)_2_, **(H)** G_1_-(DHBS)_2_, **(I)** G_2_-(DHBS)_2_, and **(J)** DHBS between five experimental groups, including UTI89 grown in low and high iron media (wild type and wild type+Fe, respectively), the enterobactin-null mutant UTI89∆*entB* in low and high iron media (*entB* and *entB* +Fe, respectively), and the enterobaftin-only mutant UTI89∆*ybtS*∆*iroA* in low iron medium (∆*ybtS*∆*iroA*). Statistics were performed using unpaired *t* test with P ≤ 0.05 considered as statistically significant. ns: not significant. *: *P* <= 0.05. **: *P* < 0.01. ***: *P* < 0.001. ****: *P* < 0.0001.

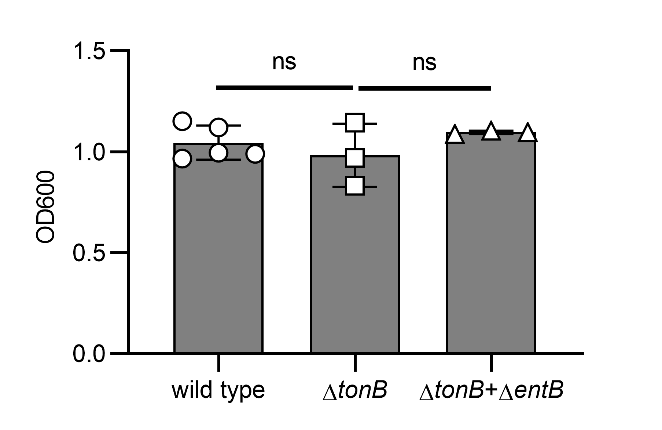

**Figure S15. Bacterial growth of wild type UTI89, import-deficient UTI89∆*tonB* mutant, and the co-culture of import-deficient UTI89∆*tonB* and biosynthesis-deficient UTI89∆*entB* mutants in iron restricted media**. wild type: UTI89. ∆*tonB*: import-deficient UTI89∆*tonB* mutant. ∆*tonB*+∆*entB*: co-culture of import-deficient UTI89∆*tonB* and biosynthesis-deficient UTI89∆*entB* mutants. Statistics were performed using 1-way ANOVA with Dunnett’s multiple-comparison test with P ≤ 0.05 considered as statistically significant. ns: not significant. *: *P* <= 0.05. **: *P* < 0.01. ***: *P* < 0.001. ****: *P* < 0.0001.

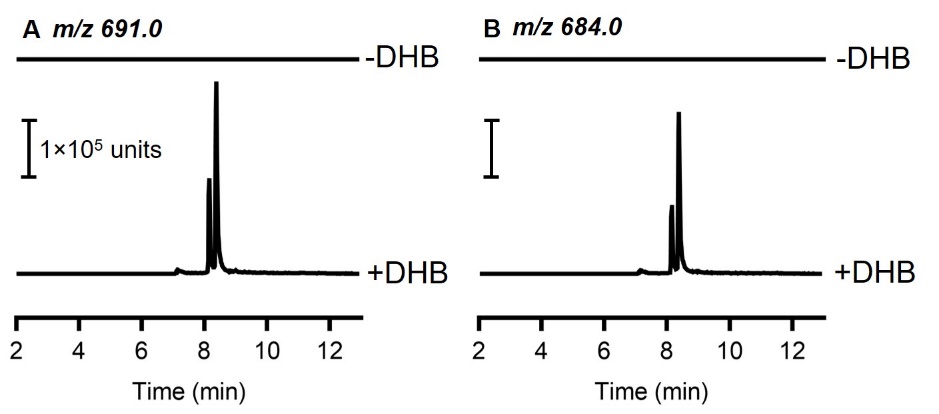

**Figure S16.** **UTI89 can use exogenous 2,3-dihydrobenzoic acid (DHB) to support enterobactin biosynthesis. (A**) LC-MS/MS detection of ^13^C_23_-substituted Ent ([M-H]^-^, *m/z* 698) in ^13^C_3_-glycerol culture medium conditioned by UTI89 grown without (-DHB) or with (+DHB) 200 µM unlabeled DHB. (**B**) LC-MS/MS detection of ^13^C_16_-substituted Ent ([M-H]^-^, *m/z* 677) in ^13^C_3_-glycerol culture medium conditioned by UTI89 grown without (-DHB) or with (+DHB) 200 µM unlabeled DHB.

**SUPPLEMENTARY TABLES**

**Table S1.** Thirty most highly upregulated metabolites in UTI89 exometabolome under iron-restricted condition.

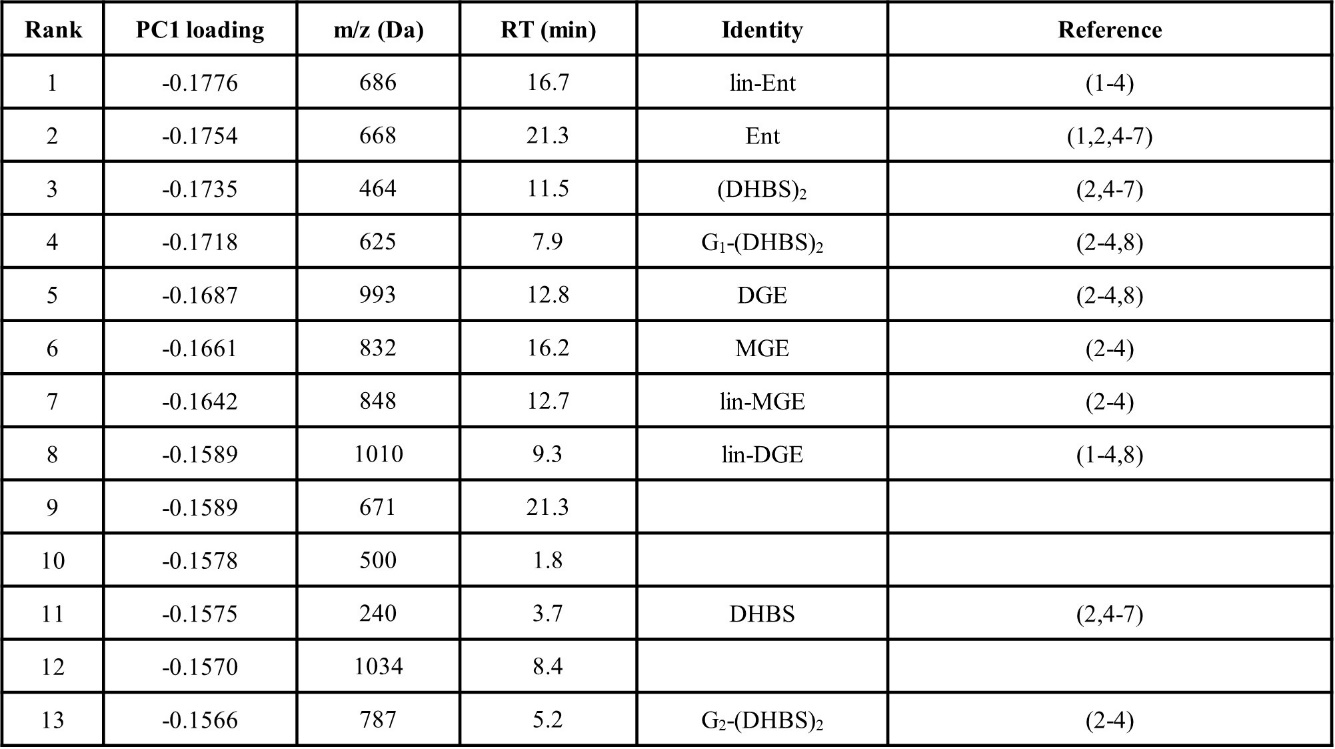

**Table S2.** Targeted LC-MS/MS methods for detecting and quantifying isotope-labelled CE produced by UTI89 grown in ^13^C_3_-glycerol M63 minimum medium with the supplement of 200 uM ^12^C-DHB.

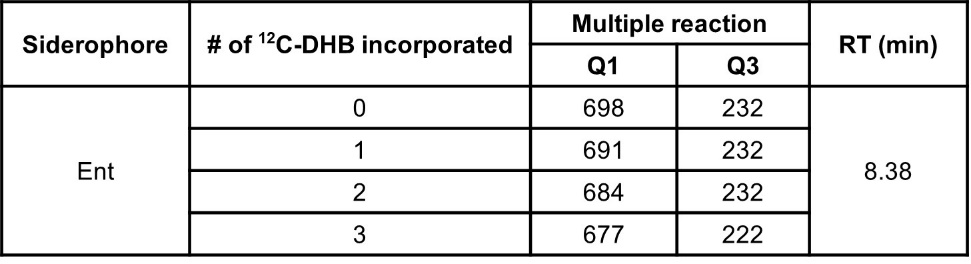

**Table S3.** Strains used in this study

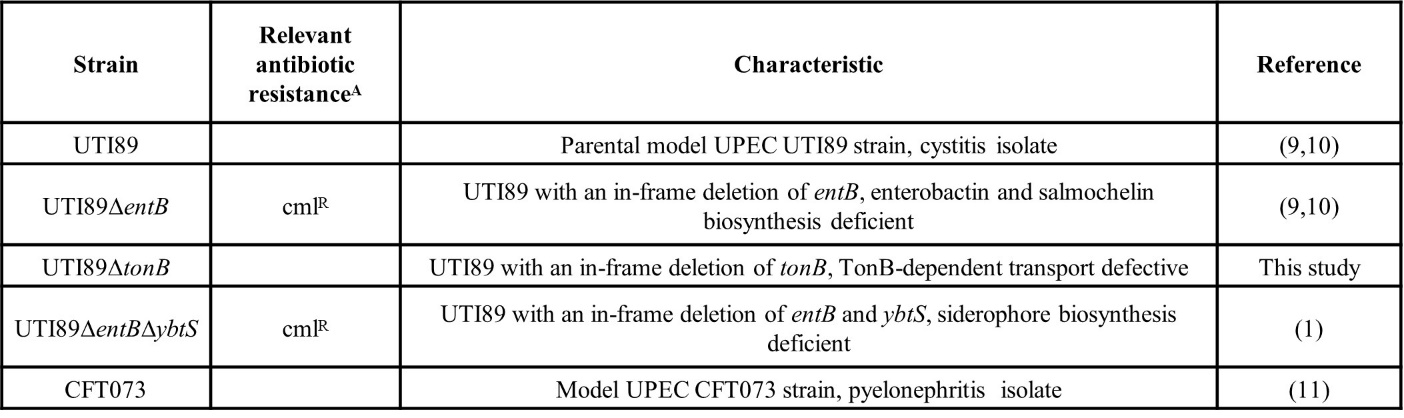

^A^ cml^R^, resistance to chloramphenicol antibiotic
